# Structural Dynamics of the Dengue Virus Non-structural 5 (NS5) Interactions with Promoter Stem Loop A (SLA)

**DOI:** 10.1101/2024.12.03.626708

**Authors:** Juliet O. Obi, Kyle C. Kihn, Linfah McQueen, James K. Fields, Greg A. Snyder, Daniel J. Deredge

**Author notes:** Corresponding author, **Correspondence:** Daniel Deredge, Department of Pharmaceutical Sciences, University of Maryland, Baltimore 20 North Pine Street, Pharmacy Hall, Room N529, Baltimore, MD 21201. Georgetown University, Washington, DC, 20057, USA. Department of Biophysics and Biophysical Chemistry, Johns Hopkins School of Medicine, Baltimore, Maryland, 21205, USA.

## Abstract

The dengue virus (DENV) NS5 protein plays a central role in dengue viral RNA synthesis which makes it an attractive target for antiviral drug development. DENV NS5 is known to interact with the stem-loop A (SLA) promoter at the 5’-untranslated region (5’-UTR) of the viral genome as a molecular recognition signature for the initiation of negative strand synthesis at the 3’ end of the viral genome. However, the conformational dynamics involved in these interactions are yet to be fully elucidated. Our study explores the structural dynamics of NS5 from DENV serotype 2 (DENV2 NS5) in complex with SLA, employing surface plasmon resonance (SPR), hydrogen - deuterium exchange coupled to mass spectrometry (HDX-MS), computational modeling, and cryoEM single particle analysis to delineate the molecular details of their interaction. Our findings indicate that DENV2 NS5 binds SLA in a closed conformation with significant interdomain cooperation between the methyltransferase (MTase) and RNA-dependent RNA polymerase (RdRp) domains, a feature integral to the interaction. Our HDX-MS studies reveal SLA-induced conformational changes in both domains of DENV2 NS5, reflecting a potential mechanism for dengue NS5’s multifunctional role in viral replication. Lastly, our cryoEM structure provides the first visualization of the DENV2 NS5-SLA complex, confirming a conserved SLA binding mode across DENV serotypes. These insights obtained from our study enhance our understanding of dengue NS5’s complex conformational landscape, supporting the potential development of antiviral strategies targeting dengue NS5’s conformational states.

## Introduction

Flaviviruses are positive-sense, single-stranded RNA viruses which give rise to many arthropod-borne viral infections globally, including the mosquito-borne dengue fever [1]. Dengue fever is caused by the dengue virus (DENV), which is primarily transmitted by *Aedes aegypti* mosquitoes, and is known to be the most prevalent mosquito-borne disease globally. Approximately 400 million people get infected with the virus annually [2], with the virus circulating mainly in tropical and sub-tropical regions, including South-East Asia, the Americas, Africa, Western Pacific, and Eastern Mediterranean regions [3]. Dengue is currently endemic in over 100 countries and affects more than 2.5 billion people living in the tropics and subtropics [4]. There are four known serotypes of the dengue virus (DENV 1-4) and humans who have been infected with one serotype, can be re-infected with another serotype leading to severe dengue disease, characterized by dengue hemorrhagic fever (DHF) and dengue shock syndrome (DSS) [5]. Severe dengue is thought to arise due to the antibody-dependent enhancement (ADE) mechanism and can be fatal if untreated [6, 7]. Although the mortality rate of severe dengue is relatively low, the disease has become a global health concern in the last few decades due to the increasing geographic expansion of *Aedes aegypti* mosquitoes which transmit the virus [8]. To date, there are no antiviral drugs approved for the treatment of dengue infection, and there are no universal vaccines available to protect people from all the dengue serotypes [9, 10].

The dengue viral replication complex (RC), which consists of some virally-encoded non-structural (NS) proteins, and the template viral RNA genome along with some host factors, is tasked with replicating the viral genome and capping the viral RNA at the 5’ untranslated region (5’-UTR) [11]. The non-structural 5 (NS5) protein is the largest and most conserved protein encoded by flaviviruses. As part of its canonical replicative function, it is a key component of the replication complex with multiple enzymatic and biological functions and is a major target for antiviral drug development [12]. The dengue virus NS5 contains an N-terminal methyltransferase (MTase) domain, with guanylyltransferase and methyltransferase activities, and a C-terminal RNA-dependent RNA polymerase (RdRp) domain, responsible for *de novo* RNA synthesis.

Dengue NS5 is also known to interact with structural elements in the 5’ and 3’ untranslated regions to promote viral RNA synthesis [11, 13–16]. The 5’-UTR in particular, contains a 70-nucleotide element known as stem loop A (SLA) which serves as a recognition motif for NS5 to discriminate the viral genome from the host mRNA. The recognition of SLA by NS5 drives genome cyclization, recruiting NS5 to the 3’ end of the viral genome for the initiation of negative-strand RNA synthesis [17–19]. Previous studies have characterized the binding of dengue NS5 to SLA with nanomolar binding affinity reported and the site of SLA binding predicted to be on the RdRp domain [20, 21]. A recent study employed cryo-EM analyses to elucidate the structure of NS5 from DENV3 in complex with SLA for initiation of replication and the NS5-NS3 complex for RNA elongation [22]. However, the conformational dynamics of the interactions between NS5 and SLA remain unclear for the different dengue serotypes. Two global conformations of full-length NS5 have been observed in flaviviruses based on the orientation of the MTase and RdRp domains connected by a flexible linker [16]. To date, all of the solved experimental structures of DENV3 NS5 are in the closed (compact) conformation, including the cryo-EM structure with SLA and NS3 [22]. The experimental structures of DENV2 NS5, however, have been observed in both the closed and open (extended) conformations [12]. Specifically, four out of five structures in the protein data bank (PDB), including a recent cryo-EM structure with human STAT2 as part of NS5’s non canonical function, are in the open conformation, with one apo crystal structure in the closed conformation [12, 16, 23]. This raises the question of whether DENV2 NS5 binds SLA in the open or closed conformation. To address this question, we have studied the interaction of NS5 with SLA in the context of serotype 2 DENV. In this study, we characterized the binding interactions of the full-length and individual domains of DENV2 NS5 with SLA using surface plasmon resonance (SPR) studies. Then, we described the conformational dynamics which contribute to the coordination of the RdRp and MTase domains during the binding interaction of full-length DENV2 NS5 to SLA using hydrogen-deuterium exchange coupled with mass spectrometry (HDX-MS). Using a combination of protein-RNA docking, molecular dynamics (MD) simulations and HDX-based ensemble reweighting (HDXer), we propose a DENV2 NS5-SLA model which conforms the most with our HDX-MS data. Finally, using cryo-EM single particle analysis, we determined the structure of DENV2 NS5 complexed with SLA for the first time. Taken together, our studies provide more intricate details into the dynamic conformational selection signature of dengue NS5 for performing its canonical and non-canonical functions.

## Materials and methods

### Stem Loop A RNA Construct

The first 70 nucleotides corresponding to SLA from the dengue virus serotype 2 genome (New Guinea C strain, GenBank no AF038403.1) was synthesized by GenScript in non-biotin and biotin-tagged forms. Lyophilized RNA samples were dissolved in respective buffers used for experiments, heated to 70°C for 2 min, and snapped cooled on ice for 1 min prior to use.

### Plasmid Construction

The cDNAs encoding full-length DENV2 NS5, RdRp and MTase domains (New Guinea C strain, GenBank no AF038403.1) were codon-optimized and cloned into the pET28a(+)-TEV vector (GenScript) with a hexahistidine tag, and a tobacco etch virus (TEV) protease cleavage site at the N-terminal region. The DNA sequences were verified by Eurofins Genomics before the plasmids were transformed into *Escherichia coli* BL21-CodonPlus (DE3)-RIL competent cells (Agilent Technologies) for protein expression.

### Protein Expression and Purification

*E. coli* BL21-CodonPlus (DE3)-RIL cells expressing full-length DENV2 NS5, RdRp or MTase domains were grown in terrific broth (TB) supplemented with 50 µg/mL kanamycin and 34 µg/mL chloramphenicol at 37°C until an OD_600_ between 0.8 and 1.0 was reached. The cells were placed on ice for 2 hours, and afterwards were induced by adding 0.25 mM isopropyl β-D-1-thiogalactopyranoside (IPTG). The induced cells were transferred to a 16°C incubator and grown overnight for 16 hours. The cells were harvested by centrifugation at 8,000 x g and resuspended in lysis buffer supplemented with EDTA-free protease inhibitor cocktail tablets (Roche), 0.1 mg/mL lysozyme and 0.1% Triton (Sigma-Aldrich). For the full-length NS5 and RdRp domain, lysis buffer containing 20 mM sodium phosphate pH 7.0, 500 mM NaCl, 50 mM L-Arginine, 50 mM L-Glutamic acid, 10 mM Imidazole, 5 mM β-mercaptoethanol (BME) and 10% glycerol was used. For the MTase domain, lysis buffer containing 50 mM HEPES pH 7.5, 500 mM NaCl, 10 mM Imidazole, 5 mM magnesium sulphate, 5 mM BME and 10% glycerol was used. The resuspended cells were homogenized in a microfluidizer (Microfluidics) and centrifuged at 10,000 rpm. The cell supernatant was loaded onto an equilibrated HisTrap HP nickel affinity column (Cytiva) and the column was connected to an AKTA pure 25 M system (Cytiva) for purification. The column was washed with wash buffer (20 mM sodium phosphate pH 7.0, 500 mM NaCl, 40 mM Imidazole, 5 mM BME and 10% glycerol for full-length NS5 and RdRp domain; 50 mM HEPES pH 7.5, 500 mM NaCl, 40 mM Imidazole, 5 mM magnesium sulphate, 5 mM BME and 10% glycerol for MTase domain) and the proteins were eluted with a linear imidazole gradient from 40 mM to 1 M with the proteins eluting at 400 mM imidazole. The fractions containing the eluted proteins were pooled and dialyzed in SEC buffer containing 20 mM HEPES pH 7.0 (or pH 7.5 for the MTase domain), 150 mM NaCl and 2 mM TCEP. The dialyzed proteins were loaded onto an equilibrated HiLoad 16/600 Superdex 200 size exclusion column (Cytiva) for SEC purification (except the full-length NS5 protein which was loaded onto a HiTrap Heparin HP column for ion-exchange purification prior to SEC purification). The proteins eluted as monomers and their purity verified by SDS-PAGE were estimated to be >95% (Supplementary Fig. 1-2). Protein concentrations were calculated from optical absorbance measurements using extinction coefficients of 217260 M^-1^cm^-1^, 172340 M^-1^cm^-1^ and 46410 M^-1^cm^-1^ for full-length NS5, RdRp and MTase domains respectively.

### Analytical Ultracentrifugation (AUC)

The homogeneity and monodispersity of DENV2 NS5 and SLA in solution were determined by analytical ultracentrifugation (AUC) sedimentation velocity (SV) studies. For DENV2 NS5, 3.7 µM samples were prepared in buffer containing 20 mM HEPES, 300 mM NaCl, 2mM TCEP pH 7.0. For SLA, 1 µM samples were prepared in buffer containing 20 mM Tris, 150 mM NaCl and 2mM MgCl_2_ pH 7.4. SV experiments were performed at 20°C by UV intensity detection using a 4-hole An-60 Ti rotor and standard 2-channel 12-mm Epon charcoal-filled centerpieces (Beckman Coulter). Samples were run at 26,000 rpm, with 400 scans collected at wavelengths monitored at 280 nm and 260 nm for DENV2 NS5 and SLA respectively. AUC data were analyzed with Sedfit 16.1c [24]. Hydrodynamic corrections for buffer density and viscosity were estimated to be 1.0122 g/mL and 1.044 cP for DENV2 NS5 (1.0339 g/mL and 1.3939 cP for SLA) using Sednterp 3.0 [25]. The partial specific volume of DENV2 NS5 and SLA were determined from their sequences using Sednterp 3.0 and the NucProt calculator [26] respectively. SV data were analyzed using the Lamm equation continuous sedimentation coefficient distribution c(s) model [24]. Results from the AUC experiments are shown in Supplementary Fig. 3.

### Electromobility Shift Assays (EMSA)

Stoichiometry of the NS5-SLA complex was analyzed on a 10% precast native polyacrylamide gel (Bio-Rad). Gels were prerun at 4°C for 90 min at 120 V in 1X TBE. Samples were prepared in binding buffer containing 20 mM Tris, 150 mM NaCl, 2 mM MgCl_2_, 10% glycerol pH 7.4 and incubated at room temperature for 30 minutes. 10 μL samples containing 50 nM 3’biotinylated SLA alone and in complex with increasing concentrations of DENV2 NS5 were loaded in each well, and were run in 1X TBE at 4°C for 2 hr at 120 V. The gel was transferred to a positively charged nylon membrane (Thermo Fisher Scientific) using a Trans-Blot^®^ SD Semi-Dry transfer cell (Bio-Rad). To covalently bind the 3’biotin-tagged SLA probe to the membrane, it was crosslinked using a UV crosslinker (Fisher Scientific) for 2 min at 120mJ. A chemiluminescence nucleic acid detection module kit (Thermo Fisher Scientific) including a streptavidin-horseradish peroxidase enzyme was used for blocking and washing the membrane. Chemiluminescence detection at a wavelength of 425 nm was performed after exposing the membrane to a Luminol/Enhancer substrate solution mixed with hydrogen peroxide for one minute using an Odyssey Fc imager (Li-Cor Biosciences). Data from the EMSA experiments is shown in Supplementary Fig. 4a.

### Dynamic Light Scattering (DLS)

DLS experiments were performed to evaluate the particle size distribution of DENV2 NS5, SLA and the NS5-SLA complex using a Zetasizer Nano S instrument (Malvern). Samples were diluted in DLS buffer containing 20 mM HEPES, 150 mM NaCl and 2 mM TCEP pH 7.0. 1 µM samples of NS5 alone, SLA alone and the complex in a 1:1 molar ratio was loaded in a ZEN2112 Quartz cuvette before measurement in the Zetasizer Nano S instrument with a backscatter detection system at 173° at 25°C. Intensity-weighted size distributions were evaluated by the Zetasizer Nano software using the non-negative least-squares algorithm for the deconvolution of correlation curves to obtain intensity-weighted size distribution data. Data from the DLS studies is shown in Supplementary Fig. 4b.

### Surface Plasmon Resonance (SPR)

The binding affinities of DENV2 NS5-SLA, RdRp-SLA, and MTase-SLA protein-RNA interactions were determined by surface plasmon resonance (SPR). SPR experiments were performed using a benchtop OpenSPR Rev4-XT (Nicoya Lifesciences). SLA and protein samples were prepared in SPR buffer containing 10 mM HEPES, 150 mM NaCl, 3 mM EDTA, 0.05% Tween 20 pH 7.4. Streptavidin (0.5 µM) was used to activate biotin sensor chips on flow cell channels 1 and 2. 50 nM SLA with a 3’biotin tag was immobilized on channel 2 only and both channels were subsequently blocked with biocytin to minimize non-specific binding. DENV2 NS5, RdRp and MTase analyte samples in increasing concentrations were flown through both channels 1 and 2 for a 200 s contact time and 450 s dissociation time. Sensorgrams were acquired for all titrated protein concentrations, and the data was analyzed to determine the association rate and dissociation rate using the TraceDrawer software (Ridgeview Instruments). Sensorgrams from all SPR experiments were referenced against the control flow cell (channel 1) and buffer blank injections. All experiments were done in triplicates. SPR data were fit to a 1:1 binding model in Tracedrawer to determine the binding affinity (K_D_) of the interactions.

### Hydrogen-Deuterium Exchange coupled to Mass Spectrometry (HDX-MS)

Undeuterated control experiments were performed for peptide identification and coverage maps for DENV2 NS5 and NS5-RdRp proteins. Undeuterated control reactions were performed as follows: 2 µL of 10 µM protein in 20 mM HEPES pH 7.4, 150 mM NaCl, 2 mM TCEP was diluted with 18 µL of ice-cold quench (100 mM Glycine pH 2.5, 6 M Urea, 500 mM NaCl, 20 mM TCEP) for 2 min prior to dilution with 150 µL of 100 mM Glycine pH 2.5. The undeuterated samples were injected into a Waters HDX system (Waters, Milford, MA) equipped with an M-class UPLC system and an in-line pepsin digestion column. The pepsin column was manually packed in house using agarose immobilized pepsin (Thermo Fisher Scientific) and column cartridges (Upchurch Scientific^®^). Peptic fragments were trapped on an Acquity UPLC BEH C18 peptide trap column and separated on an Acquity UPLC BEH C18 analytical column. A 7 min, 5% to 35% acetonitrile in 0.1% formic acid gradient was used to elute peptides directly into a Waters Synapt G2-Si mass spectrometer (Waters, Milford, MA). MSe data were acquired with a 20 to 30 V ramp trap collision energy (CE) for high energy acquisition of product ions and continuous lock mass (Leu-Enkephalin) for mass accuracy correction. Peptides were identified using the ProteinLynx Global Server 3.0.3 (Waters). Dynamx 3.0 software (Waters) was used for peptide processing with a filter of 0.3 fragments per residue applied. Sequence coverage maps obtained for DENV2 NS5 and NS5-RdRp proteins are shown in Supplementary Figures 5-6. Hydrogen-deuterium exchange reactions for DENV2 NS5 and NS5-RdRp apo proteins and the proteins in complex with SLA were carried out by manual injections. The deuteration reaction workflow for the apo proteins was as follows: 2 µL of 10 µM protein added to 2 µL H_2_O buffer (20 mM HEPES pH 7.4, 150 mM NaCl, 2 mM TCEP) was incubated in 16 µL D_2_O buffer (20 mM HEPES pD 7.4, 150 mM NaCl, 2 mM TCEP). The 20 µL reaction was quenched at various times with 80 µL of ice-cold quench (100 mM Glycine pH 2.5, 6 M Urea, 500 mM NaCl, 20 mM TCEP) with a 30 s wait time on ice prior to dilution with 150 µL of ice-cold 100 mM Glycine pH 2.5. The deuteration reaction workflow for the SLA-bound proteins was as follows: 2 µL of 10 µM protein was mixed with 2 µL of 20 µM SLA and after 2 min of equilibration, the complex was incubated in 16 µL D_2_O buffer (20 mM HEPES pD 7.4, 150 mM NaCl, 2 mM TCEP). The 20 µL reaction was quenched at various times with 80 µL of ice-cold quench (100 mM Glycine pH 2.5, 6 M Urea, 500 mM NaCl, 20 mM TCEP) with a 30 s wait time on ice prior to dilution with 150 µL of ice-cold 100 mM Glycine pH 2.5. All deuteration reactions were performed at 25°C at five reaction time points of 10 s, 1 min, 10 min, 1 hr, and 2 hr. After quenching of the apo and complex deuterated samples, the 250 µL total reaction was injected into the mass spectrometer and LC/MS acquisition was performed similarly to the undeuterated controls. The deuteration time points for all the samples were acquired in triplicate. Fully deuterated controls were performed for normalization purposes and the reaction workflow was as follows: 10 µL of 20 µM protein was incubated with 10 µL of 20 mM HEPES pH 7.4, 8 M Urea, 20 mM TCEP, and the protein was incubated overnight to unfold. 4 µL of the unfolding reaction was diluted with 16 µL of D_2_O buffer, pD 7.4, and allowed to deuterate for at least 2 hr at 25°C. The reaction was quenched with 80 µL of ice-cold quench and diluted with 150 µL of ice-cold 100 mM Glycine pH 2.5 before injection and LC-MS acquisition as described earlier. Spectral curation, centroid calculation, and deuterium uptake analysis of all identified peptides were performed using the Dynamx 3.0 software (Waters). The normalized percent deuterium uptake (%D) for each peptide, at deuterium time *t*, was calculated as described in Eq. 1 below

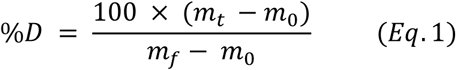

where *m_t_, m_0_,* and *m_f_* are the centroid masses at incubation time *t*, the undeuterated control, and the fully deuterated control, respectively. Bimodal deconvolution of EX1 peptides were performed using the HX-Express3 software [27, 28]. A summary of the HDX data is shown in Supplementary Table 1.

### Computational Docking and Modeling of the DENV2 NS5-SLA Complex

DENV NS5 (New Guinea C strain) was modeled with SLA in both open and closed conformations. Prior to modeling the complex, missing loop residues within the protein were filled using the SWISS-MODEL homology modeling server [29]. The User Template input option was used, with the DENV2 NS5 sequence (New Guinea C strain) uploaded as the target sequence, and apo crystal structures of DENV2 NS5 in the open and closed conformations (PDB ID: 5ZQK [16] and 6KR3 [12] respectively) uploaded as template files. Missing loop residues at the N-and C-terminals were modeled with ModLoop [30, 31]. The DENV2 NS5-SLA complex in both conformations were modeled using the HADDOCK2.4 web server [32, 33], with the apo NS5 structures described earlier and the crystal structure of DENV2 SLA with the tRNA portion removed (PDB ID: 7LYF [34]) uploaded as input structures. Protein residues on the MTase domain and RdRp thumb domain which were part of peptides observed to undergo HDX protection based on our HDX-MS data were selected as active residues for the docking process. Three stages of docking were performed including a rigid-body energy minimization, a semi-flexible refinement, and a final refinement of 200 modeled structures with short MD simulations in explicit solvent. The 200 modeled structures were scored, ranked, and clustered based on a HADDOCK score which is a weighted sum of van der Waals, electrostatic, desolvation and restraint violation energies together with buried surface area. For DENV2 NS5-SLA structures modeled in the open conformation, HADDOCK clustered 128 structures in 14 clusters. For DENV2 NS5-SLA structures in the closed conformation, HADDOCK clustered 185 structures in 18 clusters. The best 4 structures of the top 10 HADDOCK-scored clusters from each open and closed NS5-SLA conformation were downloaded from the HADDOCK website.

### Ranking of HADDOCK-generated Docking Poses using an HDX-based Scoring Function

To determine which of the top HADDOCK-scored DENV2 NS5-SLA modeled complexes agrees the most with our experimental HDX-MS data, the ‘calc-HDX’ function which is part of the HDX ensemble reweighting (HDXer) tool was used to calculate the percent deuterium uptake of the modeled structures [35, 36]. To calculate the percent deuterium uptake, the phenomenological equation [37] was used to calculate protection factors for each backbone amide hydrogen based on the Best & Vendruscolo protection factor model as described in Eq. 2 below

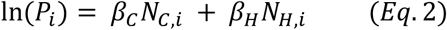

Where (*P_i_*) is the protection factor at residue *i*, *N_c,i_* is the ensemble average of the number of nonhydrogen atoms within 6.5 Å of the backbone nitrogen atom of the residue*, and N_H,i_* is the ensemble average of the number of hydrogen bonds formed by the backbone amide hydrogen of the residue. β_*C*_ and β_*H*_ are scaling factors for the backbone nitrogen atom and backbone amide hydrogen respectively. The atoms of the two neighboring residues on each side of the residue were omitted in the calculation of *N_c,i_*. Scaling factors of 0.35 and 2.0 were used for β_*C*_ and β_*H*_ respectively [37]. The computed protection factors were used to calculate peptide-level deuterium fractional uptake (*D_j,t_^sim^*) as a function of time *t* of exchange as described in Eq. 3 below

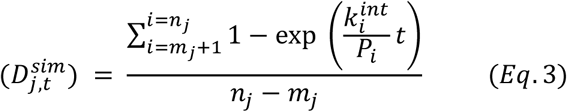

where *m_j_* and *n_j_* are the starting and ending residue numbers of the *i*^th^ peptide which were chosen to match the peptide segments in the experimental HDX-MS data. The first residue of each peptide segment, and proline residues which do not have a backbone amide hydrogen were excluded from the calculations. The intrinsic rate of exchange *k_int_* has been empirically determined by previous studies [38, 39]. The ‘protonly’ argument in the ‘calc-HDX’ function was set to false to allow any hydrogen bonding and heavy atom contacts of SLA to the DENV2 NS5 protein to be explicitly included in the protection factor calculation. The top-scored DENV2 NS5-SLA models generated by HADDOCK for both open and closed conformations were subsequently scored using an HDX-based scoring function as previously described [40]. Briefly, the HDX-based scoring function is applied by determining the root mean squared error between the experimental and calculated difference in percent deuterium uptake (Δ%D RMSE) between the apo and SLA- bound state for all observed peptides at different deuterium incubation time points. The experimental Δ%D (apo DENV2 NS5 – SLA-bound NS5) is determined by HDX-MS as described in the HDX-MS method and the computationally calculated Δ%D is determined by the ‘calc-HDX’ function as described earlier. The top 20 SLA-bound DENV2 NS5 docking poses based on their Δ%D RMSE values are shown in Table 1. The best-scored docking poses with the lowest Δ%D RMSE values in both conformations were aligned with the DENV3 NS5-SLA cryoEM structure (PDB ID 8GZP, Supplementary Fig. 7).

**Table 1.** HDX-based scoring of HADDOCK-generated DENV2 NS5-SLA docking poses. The top 20 docking poses based on their Δ%D RMSE values, and z-scores are shown for both closed and open DENV2 NS5 conformations respectively. The best scored docking poses for both closed and open conformations (cluster 4_4 and cluster 12_2 respectively) were aligned with the DENV3 NS5-SLA cryoEM structure as shown in Supplementary Fig. 7.

| Closed |  |  | Open |  |  |
| --- | --- | --- | --- | --- | --- |
| Model | $\Delta\%D$ RMSE | z-score | Model | $\Delta\%D$ RMSE | z-score |
| Cluster 4_4 | 0.1265 | -2.44 | Cluster 12_2 | 0.1402 | -2.31 |
| Cluster 11_3 | 0.1299 | -1.77 | Cluster 1_1 | 0.1406 | -2.15 |
| Cluster 8_4 | 0.1311 | -1.54 | Cluster 3_4 | 0.1437 | -0.93 |
| Cluster 2_1 | 0.1323 | -1.31 | Cluster 7_4 | 0.1437 | -0.93 |
| Cluster 8_1 | 0.1329 | -1.19 | Cluster 1_4 | 0.1440 | -0.82 |
| Cluster 1_1 | 0.1333 | -1.11 | Cluster 4_2 | 0.1440 | -0.82 |
| Cluster 11_4 | 0.1363 | -0.53 | Cluster 12_3 | 0.1443 | -0.70 |
| Cluster 4_1 | 0.1365 | -0.49 | Cluster 7_3 | 0.1446 | -0.58 |
| Cluster 1_4 | 0.1369 | -0.41 | Cluster 1_3 | 0.1447 | -0.54 |
| Cluster 8_3 | 0.1370 | -0.39 | Cluster 12_1 | 0.1448 | -0.50 |
| Cluster 2_4 | 0.1375 | -0.29 | Cluster 4_1 | 0.1448 | -0.50 |
| Cluster 1_2 | 0.1387 | -0.06 | Cluster 4_4 | 0.1448 | -0.50 |
| Cluster 6_3 | 0.1388 | -0.04 | Cluster 14_2 | 0.1449 | -0.46 |
| Cluster 8_2 | 0.1388 | -0.04 | Cluster 12_4 | 0.1450 | -0.42 |
| Cluster 4_2 | 0.1389 | -0.02 | Cluster 8_3 | 0.1451 | -0.38 |
| Cluster 4_3 | 0.1392 | 0.04 | Cluster 8_2 | 0.1452 | -0.34 |
| Cluster 1_3 | 0.1394 | 0.08 | Cluster 7_1 | 0.1453 | -0.30 |
| Cluster 6_1 | 0.1394 | 0.08 | Cluster 2_3 | 0.1455 | -0.23 |
| Cluster 14_2 | 0.1395 | 0.10 | Cluster 3_2 | 0.1460 | -0.03 |
| Cluster 11_1 | 0.1397 | 0.14 | Cluster 9_2 | 0.1460 | -0.03 |

### Molecular Dynamics (MD) Simulations and HDX Maximal Entropy Reweighting (HDXer)

MD simulations of apo and docked SLA-bound DENV2 NS5 in both open and closed conformations were performed using OpenMM v8.1 [41]. The best-scored poses for SLA-bound DENV2 NS5 in both conformations were used for the simulations. Input files for the simulations including the parametrization of the SAM ligand were generated using the CHARMM graphical user interface (CHARMM-GUI) input generator web application [42]. Simulations were performed using the CHARMM36m additive force field [43] and the TIP3P water model [44]. The system was solvated in a periodic water box containing 0.15 M KCl with 1 nm box boundaries to the solute atoms. A force switching function that ranged from 1.0 to 1.2 nm was applied for the Lennard-Jones interaction calculations. Long-range electrostatic interactions were calculated using a particle mesh Ewald approach, with an Ewald error tolerance of 0.0005. A 2 fs time step was utilized for integration with temperature and pressure held constant at 298.15 K and 1 atm, respectively. A Langevin thermostat was used to maintain the temperature at 298.15 K with a frictional coefficient of 1 ps^-1^. A Monte Carlo (MC) barostat was used to hold the pressure isotopically constant at 1 bar with a pressure coupling frequency of 2 ps. The system energy was minimized before the production run using the Limited-memory Broyden-Fletcher-Goldfarb-Shanno algorithm (L-BGFS) method, in which 5000 minimization steps were performed, utilizing a convergence tolerance of 100 kJ/mol. The system was equilibrated for 125 ps in the NVT ensemble with a 1 fs time step. Positional restraints were applied to the protein’s backbone and side chain atoms during energy minimization and equilibration runs, with a force constant of 400 and 40 kJ/mol/A^2^, respectively. For the production runs, each open and closed conformations of apo and SLA-bound DENV2 NS5 was simulated for 250 ns, with structural coordinates written to the trajectory every 20 ps of simulation time, resulting in 12,500 frames total for each simulation. For the purposes of ensemble reweighting, all peptides displaying the characteristic bimodal behavior of the EX1 kinetic regime were excluded because the calculation of percent deuterium uptake from computational ensembles and reweighting are only applicable to the EX2 kinetic exchange regime of hydrogen-deuterium exchange as previously described [35, 45, 46]. For the resulting set of peptide segments, hydrogen-deuterium exchange ensemble reweighting (HDXer) was performed. A mixed ensemble of conformations was first generated by combining the open and closed MD trajectories together for both apo and SLA-bound DENV2 NS5 states respectively. The averaged peptide segment deuteration fractions over the resulting mixed ensembles were calculated using the ‘calc_HDX’ function of HDXer. Reweighting was subsequently performed using HDXer as previously described [35, 36, 40, 47]. Briefly, HDXer was utilized to adjust the weight of individual frames such that the frames which conform the most with the solution-based HDX-MS data are given more weight (upweighted). The mixed ensemble for apo DENV2 NS5 was reweighted against the target apo HDX-MS experimental data, and the mixed ensemble for SLA-bound DENV2 NS5 was reweighted against the target SLA-bound HDX-MS experimental data respectively. A summary of the MD simulations and HDXer workflow is shown in Supplementary Fig. 8.

### CryoEM Sample Preparation and Data Collection

DENV2 NS5 was mixed with SLA in a 1:2 molar ratio in buffer containing 20 mM HEPES pH 8.0, 150 mM NaCl, 2 mM MgCl_2_, and 2 mM TCEP. The DENV2 NS5-SLA complex was incubated at room temperature for 30 min, after which glutaraldehyde was added to a final concentration of 0.01%. The crosslinked complex was further incubated on ice for 1 hr, and 1 M Tris-HCl pH 8.0 was added afterwards for glutaraldehyde inactivation. The mixture was purified using size-exclusion chromatography on a Superdex 200 Increase 10/300 GL column (Cytiva) pre-equilibrated with 20 mM HEPES pH 8.0, 150 mM NaCl, 2 mM MgCl_2_, and 2 mM TCEP. Fractions corresponding to the DENV2 NS5-SLA complex were pooled and concentrated to ∼0.5 mg/mL for subsequent use. For grid preparation, 3 μL of the sample was applied to a glow-discharged Quantifoil R 1.2/1.3 mesh Cu300 holey carbon grid. The grid was blotted for 3 s, at 4°C and 100% humidity, and plunge-frozen in liquid ethane cooled by liquid nitrogen with a Vitrobot IV (Thermo Fisher Scientific). The frozen grid was loaded onto a Talos Arctica cryo-transmission electron microscope (Thermo Fisher Scientific) operating at 200 kV and equipped with a K3 direct electron detector (Gatan). A true magnification of 69,589x was used to record movies, at a pixel size of 0.718 Å. The micrographs were recorded at 41 frames, with a dose per frame rate of 1.124 e/Å^2^ and exposure time of 4.5 s, resulting in a total dose of 46.077 e/Å^2^ and a total of 3,257 micrographs collected. A summary of the data collection parameters is shown in Supplementary Table 2.

### CryoEM Data Processing

The collected cryoEM data were processed using cryoSPARC (v4.4.1) [48]. Full-frame motion correction was used for motion correction and dose-weighting of the imported movies. Contrast transfer function (CTF) estimation was done on the imported micrographs, with a pixel size of 0.718 Å. The blob picker method in cryoSPARC was used for initial particle picking, and 5,076,473 particles were extracted from 3,257 micrographs for 2D classification. Templates were selected from the initial 2D classification for another particle picking using the template picker method. 3,683,950 particles were used for subsequent 2D classifications after template picking. After multiple rounds of 2D classifications to remove bad particles, two batches of 2D classifications were used for 3D reconstruction. The first batch with 23 classes comprising of 41,889 particles were selected for 3D *ab initio* reconstruction using 3 classes of initial models. The second batch with 62 classes comprising of 94,901 particles were selected for 3D *ab initio* reconstruction using 4 classes of initial models. Homogeneous refinement was subsequently performed on the classes corresponding to the right NS5-SLA density for each batch (17,166 and 23,294 particles respectively), with no symmetry (C1) imposed. A final round of homogeneous refinement was performed by combining the particles from the previous refinements to obtain the final DENV2 NS5-SLA density map with a total of 32,126 particles. The Fourier shell correlation (FSC) criteria at 0.143 and 0.5 were used to determine the average resolution and local resolution estimation respectively [49]. The final density map was further sharpened using the automated sharpening tool in Phenix [50]. A summary of the data processing flowchart is shown in Supplementary Fig. 9.

### Model Building and Refinement

For cryoEM model building, the best-scored DENV2 NS5-SLA docking pose in the closed conformation which significantly agreed the most with our experimental HDX-MS data as described earlier was used as the initial model. Initial fitting of the initial model to the map as a rigid body was done in UCSF Chimera [51]. Afterwards, flexible fitting of the initial model to the map was performed using the Namdinator web-based tool with the default settings [52]. The DENV2 NS5-SLA atomic model was built with Coot [53] and multiple rounds of refinement were performed with the real-space refinement tool in Phenix [54]. Self-restraints were applied in the refinement process to maintain the correct base-pair geometry in the SLA portion of the complex. The refinement statistics of the final model are presented in Supplementary Table 2. Final figures were prepared with UCSF ChimeraX [55]. The model statistics were validated using MolProbity [56].

## Results

### DENV2 NS5 Binding to Stem Loop A (SLA)

Dengue NS5 specifically recognizes SLA for the initiation of negative strand RNA synthesis at the 3’ end of the dengue viral genome. Dengue NS5 is also known to recognize SLA during positive-strand RNA synthesis to form a ^m7^GpppA^m^ type 1 cap for a fully synthesized viral genome [34]. Previous studies of DENV2 NS5 with SLA had suggested that NS5-SLA interactions are primarily mediated by the RdRp domain, and that there is no significant contribution to the interaction by the MTase domain [20]. One study of DENV3 NS5 with SLA had initially suggested that SLA interacts with NS5 through the thumb subdomain of the RdRp [21], but another recent study showed that mutation of certain residues in both the MTase and RdRp domains affected the binding of NS5 to SLA [34]. More recently, cryoEM single particle analysis studies of DENV3 NS5 have revealed and confirmed that SLA binds NS5 through both MTase and RdRp domains [22]. To fully characterize the binding contributions of the individual MTase and RdRp domains to SLA, compared to full-length NS5, we conducted surface plasmon resonance (SPR) studies to determine their respective binding affinities (K_D_). We determined that the MTase domain is capable of independently binding SLA with the RdRp domain displaying a higher affinity. Indeed, the K_D_ values of the individual MTase and RdRp domains were 2 ± 0.2 µM and 143 ± 42.8 nM respectively. However, the presence of both domains was observed to increase the affinity to SLA 10-fold with a K_D_ value for full-length DENV2 NS5 of 14 ± 2.5 nM (Fig. 1b-d). To further confirm the stoichiometry and homogeneity of the NS5-SLA complex, we conducted electromobility shift assays (EMSAs) and dynamic light scattering (DLS) studies, which displayed a 1:1 stoichiometry and a homogenous particle-sized complex (Supplementary Fig. 4).

**Figure 1.**
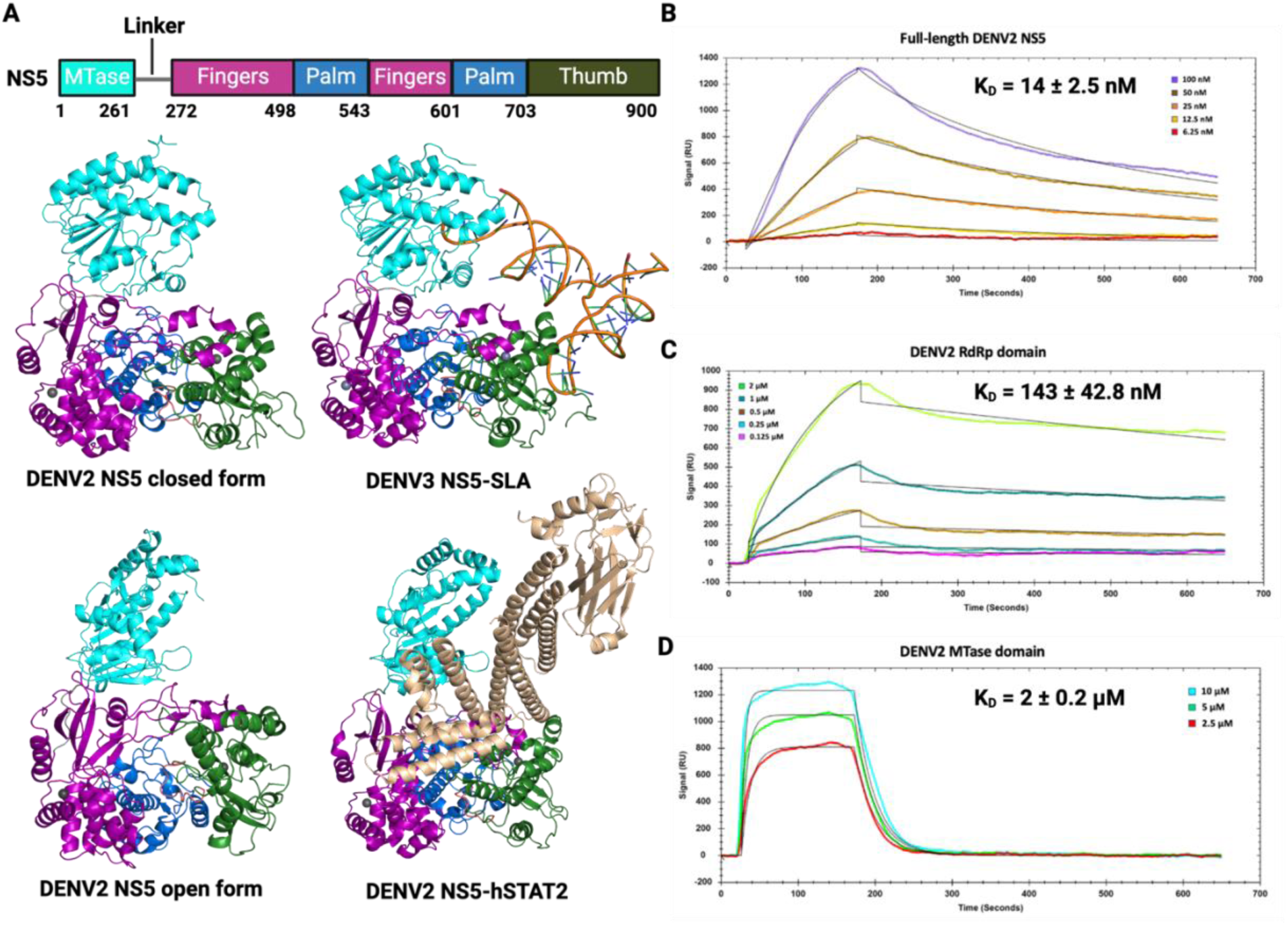
Binding Affinity Analysis of Full-length DENV2 NS5 and its Individual Domains to SLA70. **A)** DENV2 NS5 apo structures in the closed and open forms (top left and bottom left, PDB IDs 6KR3 and 5ZQK respectively), DENV3 NS5 bound to SLA (top right, PDB ID: 8GZP), and DENV2 NS5 bound to hSTAT2 in its open form (bottom right, PDB ID: 8T12). DENV2 NS5 sub-domains are colored as follows: MTase (cyan), fingers (purple), palm (marine), and thumb (forest green). Surface Plasmon Resonance (SPR) binding kinetic traces of the interactions of **B)** full-length DENV2 NS5, **C)** DENV2 RdRp domain and **D)** DENV2 MTase with SLA70. Individual traces show association and dissociation curves of the binding to SLA70 with calculated binding affinities shown.

### Mapping the Binding Interface of SLA on DENV2 NS5 with HDX-MS

Results from our SPR studies showed that both MTase and RdRp domains are also involved in SLA binding in DENV2 NS5. This led us to probe the site(s) of SLA interaction on DENV2 NS5 and delineate the conformational dynamics of apo and SLA-bound DENV2 NS5 in solution. To achieve this, we conducted HDX-MS studies on DENV2 NS5 and NS5-RdRp in their apo and SLA-bound states. For HDX-MS studies on DENV2 NS5 with SLA, the significantly protected or deprotected peptides upon SLA binding were mapped onto a DENV2 NS5 apo crystal structure for all deuteration time points (PDB ID: 5ZQK, Fig. 2a). Peptides observed to undergo significant protection from hydrogen-deuterium exchange upon SLA binding were mapped in blue color, while peptides observed to undergo deprotection from hydrogen-deuterium exchange upon SLA binding were mapped in red color. At the 10 s time point, protected peptides were observed on the MTase domain, the fingers extensions and the thumb subdomain of the RdRp (blue colored regions). At the 1 min and 10 min time points, more peptides were observed to be protected in the same domains as seen for the 10 secs time point. Interestingly, at the 10 min time point, deprotection was observed in the fingers and palm subdomain of the RdRp (red colored regions). More peptides in these regions of the RdRp domain were deprotected at the 1 hr to 2 hr time point, in addition to increasing protection seen on additional peptides on the MTase and RdRp thumb domains. We generated a difference plot showing the difference in percent deuteration incorporation between apo and SLA-bound states of DENV2 NS5 (Δ%D). The Δ%D values were plotted for each peptide fragment from the N- to the C- terminal and for each deuteration time point (Fig. 2b). We compared the difference in percent deuterium incorporation between DENV2 NS5 and NS5-RdRp in their apo and SLA-bound states by generating a difference plot for the apo and SLA-bound state of NS5-RdRp (Supplementary Fig. 10). For the NS5-RdRp domain, the peptides undergoing significant protection and deprotection upon SLA-binding were overall similar when compared to the RdRp domain in the context of full-length DENV2 NS5. However, the Δ%D values were generally lower in the context of the NS5-RdRp domain alone (Supplementary Fig. 10). To determine whether the regions of protection in our HDX-MS experiments were in agreement with the NS5-SLA binding interface in the DENV3 NS5-SLA cryoEM structure (PDB ID: 8GZP), we mapped these regions onto the corresponding peptides on DENV3 NS5. We also mapped the regions of deprotection on the DENV3 NS5-SLA cryoEM structure to show that these regions are similar to those seen in DENV2 NS5 (Fig. 2c).

**Figure 2.**
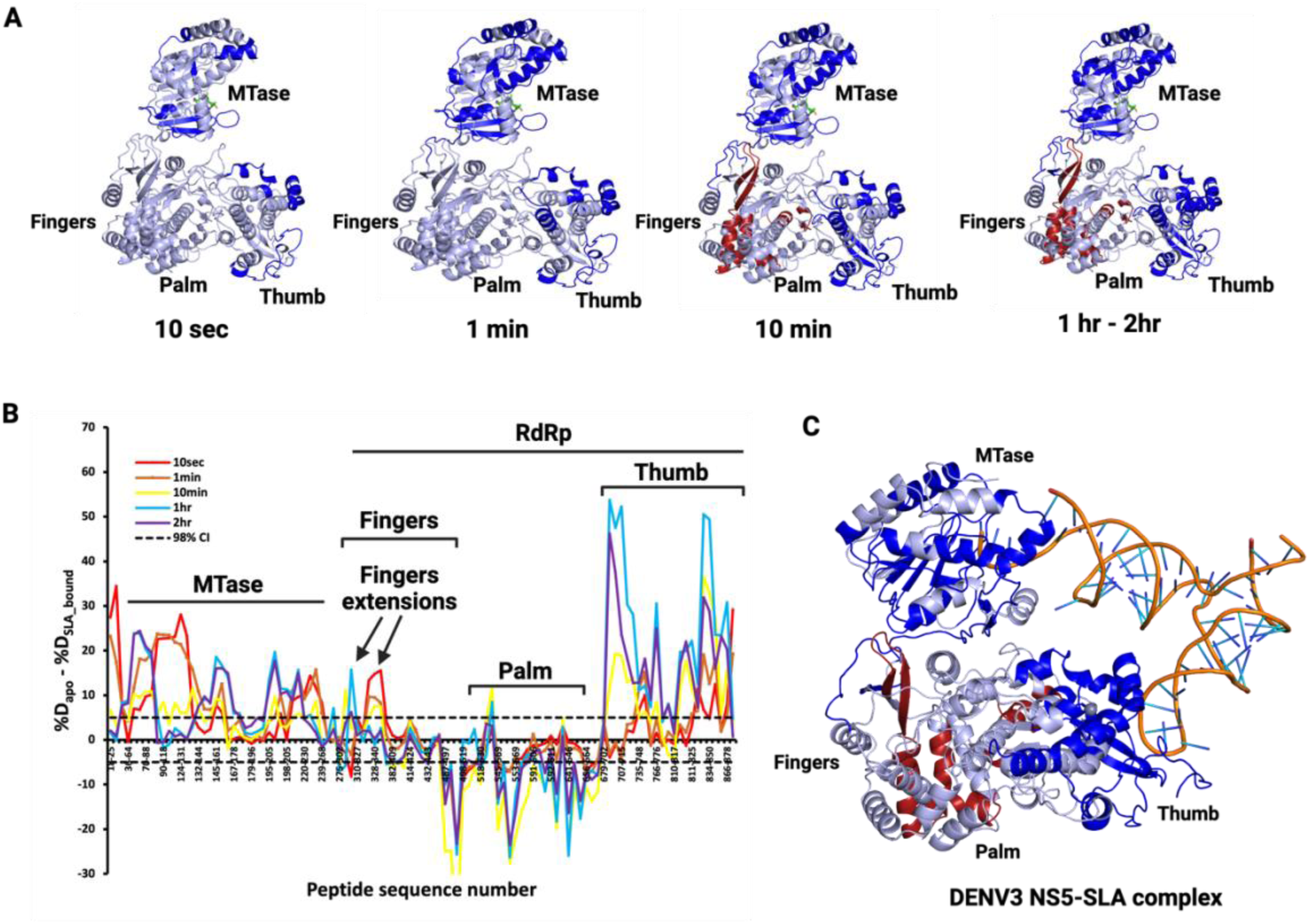
Analysis of the DENV2 NS5-SLA Binding Interface with HDX-MS. **A)** Peptic peptides with significant protection (blue-colored regions) and deprotection (red-colored regions) from deuterium labelling upon SLA binding were mapped onto the DENV2 NS5 apo crystal structure (PDB ID: 5ZQK). The significantly protected and deprotected regions throughout the deuteration time course from 10 sec to 2 hr are shown. **B)** A difference plot highlighting the differences in percent deuterium incorporation of apo *versus* SLA-bound DENV2 NS5 is shown. Peptide fragments from N- to C- terminal of the full-length DENV2 NS5 are shown from left to right based on the residue number of the first amino acid of each peptide. The difference in percent deuterium incorporation between the apo and SLA-bound states (Δ%D) is plotted for each peptide fragment and for each deuteration time point. The color coding used is based on the deuteration time points probed. The horizontal dashed lines indicate 98% confidence intervals and the different sub-domains throughout the protein are indicated. **C)** The significantly protected and deprotected regions at the 2 hr time point were mapped onto the corresponding regions on the DENV3 NS5- SLA cryoEM structure (PDB ID: 8GZP).

### Conformational Dynamics of DENV2 NS5-SLA Interactions

In HDX-MS, two broad kinetic regimes define the relative rates of local conformational dynamics with respect to the rate of hydrogen to deuterium chemical exchange. Most commonly observed under physiological conditions is the EX2 kinetic regime, wherein local regions of a protein undergo local structural fluctuations in solution which are faster than the chemical rate of exchange (ms timescale). This is observed as unimodal peptide mass spectra envelopes which gradually shift in mass over the course of the deuteration [57, 58]. EX1 kinetics, on the other hand, is thought to reflect slower conformational fluctuations typically associated with large cooperative conformational transitions. It is observed as a bimodal isotopic distribution on peptide mass spectra where the undeuterated species is representative of the population yet to undergo the correlated opening motion, and the highly deuterated species represents the population which has undergone an opening motion, with most of the amides fully exchanged [57, 59, 60]. Our HDX-MS experiments revealed bimodal isotopic envelops characteristic of EX1 exchange kinetics occurring in several regions which were seen to be responsive to SLA binding. This includes some regions in the MTase domain and RdRp fingers extensions shown to be protected from exchange upon SLA binding, and some deprotected peptide regions in the middle and pinky fingers subdomain of the RdRp domain (Fig. 3). The binding of SLA to DENV2 NS5 had varying effects on the EX1 exchange dynamics in these different peptide regions. For instance, bimodal isotopic envelops were observed in peptides 65-88 and 145-167 in the MTase domain as early as 10 s in the apo state and persisted over the deuteration time course, up to 2 hr (Fig. 3a, left and right panels). The presence of SLA however, suppressed this cooperative transition, with the bimodal isotopic distribution showing up at 60 secs, and a more significant suppression in the bimodal behavior occurring in peptide 65-88 compared to peptide 145-167. Interestingly, no bimodality was observed in the apo state for protected peptide 89-118, but it was seen at 10 secs and 60 secs in the presence of SLA (Fig. 3a, middle panel). These peptide regions when localized on the apo DENV2 NS5 crystal structure (PDB ID: 5ZQK), surround the catalytic pocket of the MTase domain where capping occurs and the S-adenosyl methionine (SAM) co-factor binds (Fig. 3b). In the RdRp domain, some minor EX1 characteristic bimodality was observed in peptide 328-341 which localizes to the index finger at 10 s in the apo state (Fig. 3d). However, the presence of SLA resulted in a mild slowdown of the EX1 kinetics, with the bimodality observed at both 10 s and 60 s deuteration time points (Fig. 3c, left panel). In addition, deprotected peptides 545-569 and 570-590 in the middle and pinky fingers respectively (Fig. 3d) were observed to undergo EX1 kinetics in the apo state. The presence of SLA however, enhanced the EX1 kinetics in these regions, with a faster rate of appearance of the highly deuterated species up to 2 hours of labeling time (Fig. 3c, middle and right panels). The plots showing the relative appearance of the highly deuterated species as a function of time for all the EX1 peptides in the MTase and RdRp domains in their apo and SLA-bound state are shown in Supplementary Fig. 11. To determine whether the presence of the MTase domain is required for the EX1 cooperative unfolding observed in the RdRp domain of DENV2 NS5, we probed the existence of bimodality in the HDX-MS experiments with NS5-RdRp and SLA. We did not observe any bimodal behavior indicative of EX1 kinetics in peptides 328–341 and 545–569 for the NS5-RdRp domain alone in both apo and SLA-bound states (Supplementary Fig. 12-13). For peptide 570–590, we observed EX1 kinetics similar to those observed in DENV2 NS5. However, an increase in the rate of appearance of the highly deuterated species was observed for NS5-RdRp in the SLA-bound state compared to DENV2 NS5 in the SLA-bound state (Supplementary Fig. 14).

**Figure 3.**
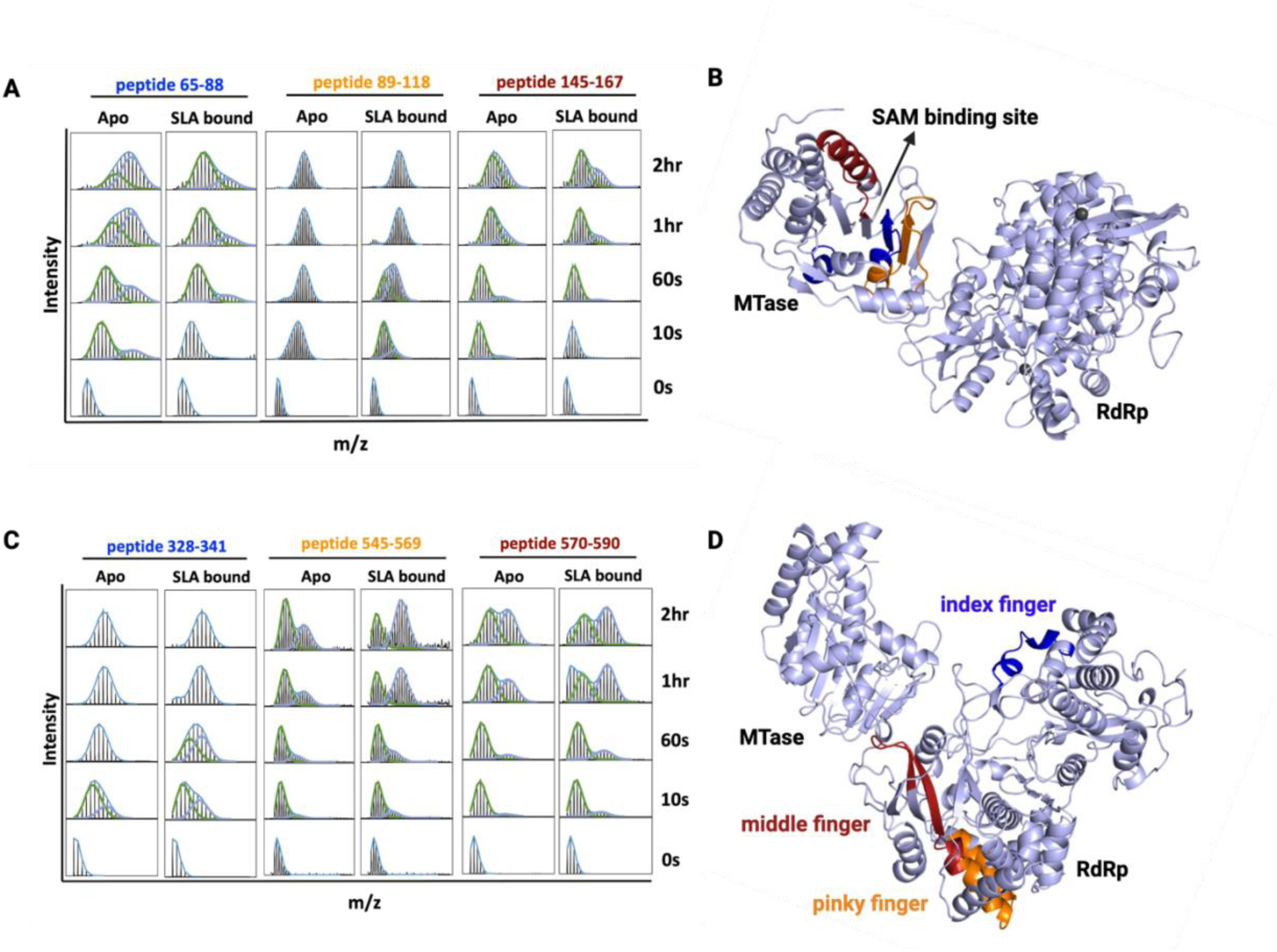
Deconvolution of Bimodal Isotopic Envelopes for DENV2 NS5 Peptides showing bimodal EX1 kinetics of deuterium exchange. **A)** The deconvoluted bimodal isotopic envelopes for EX1 peptides in the MTase domain in the apo and SLA-bound form are shown. **B)** The EX1 peptides in the MTase domain localize to the region surrounding the SAM cofactor binding site when mapped onto the DENV2 NS5 apo structure (peptide 65-88, blue; peptide 89- 118, orange; peptide 145-167, red). **C)** The deconvoluted bimodal isotopic envelopes for EX1 peptides in the RdRp domain in the apo and SLA-bound form are shown. **D)** The EX1 peptides in the RdRp domain localize to the index finger (blue), pinky finger (orange) and middle finger (red) respectively, when mapped onto the DENV2 NS5 apo crystal structure. PDB ID: 5ZQK.

### Modeling of the DENV2 NS5-SLA Complex and HDX-based Refinement of Docking Poses

More of the apo structures of DENV2 NS5 in the protein data bank were solved in the open conformation (PDB IDs 5ZQK and 6KR2), except one apo structure solved in the closed conformation (PDB ID 6KR3). Since both global conformations have been observed for DENV2 NS5, it has been suggested that they both co-exist in solution [12, 14, 16]. To determine whether the DENV2 NS5 strain we used in this study exists in the open or closed state in the apo form, and whether it binds SLA in the open or closed state, we modelled the DENV2 NS5-SLA complex in both open and closed conformations using HADDOCK. As described earlier, the best 4 structures of the top 10 HADDOCK-scored clusters from each open and closed NS5-SLA conformation were used for subsequent HDX-based refinement leading to a total of 40 docking poses for each open and closed conformation. As done in previous studies [40], we used the ‘calc-HDX’ function in the HDXer tool to apply an HDX-based scoring function, ranking the best docking models using the root mean square error (RMSE) in the difference in deuterium exchange between the computed deuterium uptake and the experimental deuterium uptake (Δ%D RMSE). The best scored models from the closed and open conformation have Δ%D RMSE values of 0.1265 and 0.1402 respectively (Table 1, Supplementary Fig. 7). Optimally, the ‘calc_HDX’ function is used to calculate predicted HDX-MS protection factors and deuterated fractions of an ensemble from molecular dynamics trajectories, and not from single protein structures. Therefore, to further confirm and validate the results from our Δ%D RMSE calculations, we employed an established strategy of HDX ensemble reweighting (HDXer) [35, 36]. We performed 250 ns conventional molecular dynamics (MD) simulations of apo DENV2 NS5 in its closed and open conformations (PDB IDs 6KR3 and 5ZQK respectively), and the best-scored DENV2 NS5-SLA docking models in both open and closed conformations based on the Δ%D RMSE values stated previously. For apo and SLA-bound states, the resulting trajectories from the open and closed conformations were combined and reweighted with HDXer using the apo and SLA-bound HDX-MS data respectively. To evaluate the conformational exploration of both open and closed states after MD simulations and HDXer, we chose the radius of gyration and the distance between residues S23 and T854 on the MTase and RdRp domains, respectively, as collective variables for a kernel density (KDEL) plot (Fig. 4). Prior to reweighting for both apo and SLA-bound states, density for the populations corresponding to both open and closed conformations were observable in the KDEL plots (Fig. 4a and 4b, top left and bottom left panels), albeit with the open conformation populations located in regions with lower density compared to the closed conformation populations. After reweighting, the weight distribution is shifted and the populations corresponding to the closed conformation populate most of the weight for both apo and SLA- bound states (Fig. 4a and 4b, top right and bottom right panels), indicating that these populations conform better with the respective solution-based HDX-MS data. Collectively, our HDXer results suggests that DENV2 NS5 adopts a closed conformation in solution, contrary to the majority of structures of DENV2 NS5 solved to date, and that SLA binding favors the closed conformation.

**Figure 4.**
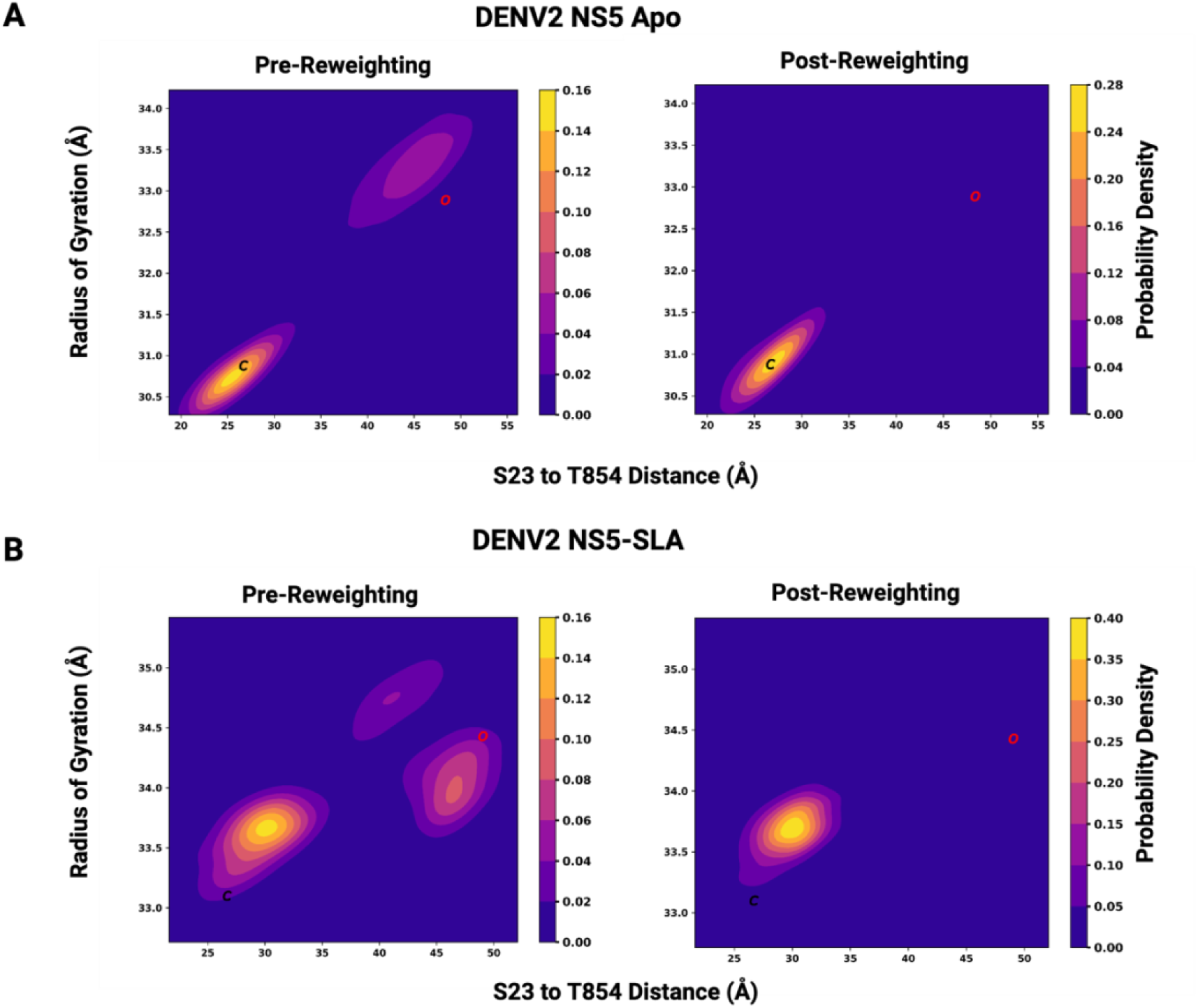
HDX Ensemble Reweighting (HDXer) of the DENV2 NS5 Apo and SLA-bound Molecular Dynamics Simulations. KDEL plots showing pre-reweighting (top left and bottom left) and post-reweighting (top right and bottom right) probability densities of the combined MD trajectories for apo DENV2 NS5 and SLA-bound DENV2 NS5 respectively. The radius of gyration (Rg) and the distance between residue S23 on the MTase domain and residue T854 on the RdRp thumb domain were used as the collective variables. The coordinates of the initial starting structures used for the MD simulations are indicated as C (black) and O (red) for the closed and open conformations respectively. For DENV2 NS5 bound to SLA, the initial starting structures for the closed and open conformations represent the best-scored docking poses based on the Δ%D RMSE values.

### CryoEM Structure of the DENV2 NS5-SLA complex

Based on our computational modeling and HDXer results, we sought to experimentally confirm whether DENV2 NS5 indeed binds to SLA in the closed conformation. To this end, we employed cryoEM single particle analysis to solve the structure of the DENV2 NS5-SLA complex. We obtained a 3.59 Å resolution map (Fig. 5a, EMD ID: 47165) and used the best-scored docking pose from the closed conformation as the initial model in building the final, deposited model (Fig. 5b, PDB ID: 9DTT). Similar to the published DENV3 NS5-SLA cryoEM structure, SLA is bound to DENV2 NS5 in its closed conformation and it adopts a V-shaped structure with the bottom stem loop and the top stem loop directly interacting with the MTase and RdRp thumb domains respectively (Fig. 5b). We aligned our final model of the DENV2 NS5-SLA complex to the DENV3 NS5-SLA cryoEM structure (PDB ID: 8GZP) and obtained an overall RMSD of 2.16 Å (Supplemental Fig. 15a) [55]. When comparing SLA in our cryoEM structure to that of the 8GZP cryoEM structure, we observe differences in their angles and junction as was reported previously [22] [34]. We also observed that most of the structural differences in SLA for both structures occur at the bottom stem-loop (Supplementary Fig. 15b). In the 8GZP cryoEM structure, the capped 5’ end of SLA extends into the catalytic binding site of the MTase domain. In our DENV2 NS5-SLA structure, we did not observe this most likely because we used an uncapped SLA for our cryoEM studies, and we were unable to model the first three nucleotides at the 5’ end. Although we obtained a cryoEM map with an overall resolution of 3.59 Å, the local resolution in various regions of the map made it difficult to properly fit some regions in our model to the map (Fig. 5c), therefore we are unable to make any inferences on any residue-specific differences in the interactions between NS5 and SLA for the DENV2 and DENV3 NS5 serotypes. However, our cryoEM results suggest that SLA interacts with NS5 in a conserved manner among the different dengue serotypes, and we are able to show, for the first time, that DENV2 NS5 binds SLA in the closed conformation.

**Figure 5.**
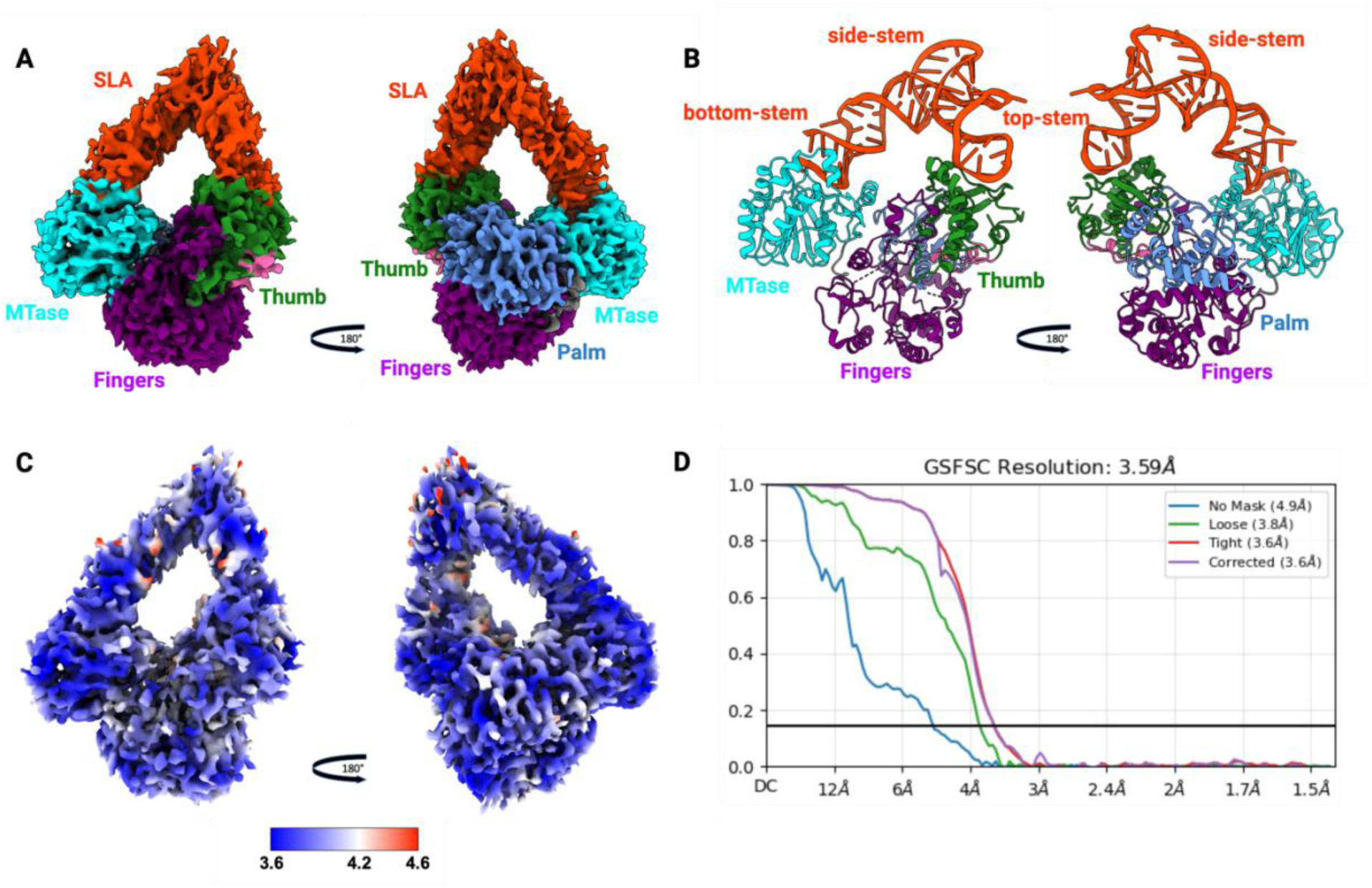
CryoEM Structure of the DENV2 NS5-SLA complex. **A)** Cryo-EM map of the complex **B)** Final model from the cryo-EM map. Both the map and the model are colored based on the different domains as follows: MTase (cyan), Fingers (purple), Palm (marine blue), Thumb (forest green). SLA is colored in red. **C)** Local resolution map of the DENV2 NS5-SLA complex **D)** Fourier shell correlation (FSC) curve of the density map of the DENV2 NS5-SLA complex based on the 0.143 cut-off in cryoSPARC.

## Discussion

Flavivirus NS5 has been shown to adopt multiple conformations in solution, which range from closed to more open conformations [14]. Studies have shown that these closed and open conformations are conserved in flaviviruses and are important for viral proliferation [12]. The conformational landscape of DENV NS5 across the dengue serotypes is complex, with the closed conformation observed in DENV3 NS5 [1], and both closed and open conformations observed in DENV2 NS5 [12, 16]. Importantly, these conformations are thought to play an important role in the recognition of binding partners for its canonical RNA replication function and non-canonical functions. For instance, DENV3 NS5’s interaction with viral promoter SLA at the 5’ UTR has been shown to be in the closed conformation [22] whereas DENV2 NS5 has been reported to interact with viral host protein hSTAT2 in the open conformation [23] (Fig. 1a). In our study, we probed the structure and dynamics of the interaction of DENV2 NS5 with SLA. Our SPR results showed that the MTase and RdRp domains can interact individually with SLA, however, cooperativity is essential between these two domains, stabilizing the interactions of full-length NS5 with SLA. This was confirmed by our HDX-MS studies which revealed protection in the MTase and RdRp domains consistent with the cryo-EM structure of DENV3 NS5-SLA [22]. Moreover, our use of integrative HDX approaches to model DENV2 NS5 indicates that, in both apo and SLA-bound form, DENV2 NS5 is more consistent with the closed conformation; an observation confirmed by cryoEM for the DENV2 NS5-SLA bound complex. The emerging picture suggests that, although several crystal structures of DENV2 NS5 have been solved in the open conformation, apo DENV2 NS5 adopts mainly a closed conformation. Such closed conformation is the conformation that favors SLA binding, and this binding mechanism is most likely conserved among the various serotypes.

More detailed analysis of our HDX-MS studies have also provided more insights into the conformational dynamics associated with the binding of SLA with dengue NS5. Indeed, the slow conformational dynamics observed as EX1 exchange kinetics in our HDX-MS studies suggest that the binding of SLA to NS5 may involve conformational rearrangements which may be reflective of both the canonical and non-canonical functions of dengue NS5. For instance, in the MTase domain, the regions where EX1 exchange kinetics was observed surround the SAM cofactor and GTP binding site. These conformational rearrangements in the MTase domain may therefore reflect important fluctuations for recognition of the capping status of the 5’ end of the genome, catalytic activity and 5’ capping of the nascent positive strand genomic RNA, or coordination of capping and polymerase activities of dengue NS5. However, in addition to the canonical replication functions of dengue NS5, this protein is also known to translocate to the nucleus to suppress human JAK-STAT signaling for counteracting the host’s immune response. For instance, studies have shown that dengue NS5 can modulate NF-κB activation by tagging the protein ERC1 for proteasomal degradation, and this mechanism has been shown to involve the MTase domain [61]. Additionally, flaviviral NS5 including DENV and Zika NS5 have been shown to suppress STAT2 to interfere with IFN signaling pathway and establish viral infection. Specifically, hSTAT2 is known to first bind in between the interdomain cleft of dengue NS5 MTase and RdRp domains, adjacent to peptides undergoing EX1 kinetics in MTase, and then a secondary interaction is established between the RdRp domain and the N-terminal domain of hSTAT2 [23, 62]. As such, it is also possible that the conformational rearrangements observed by HDX-MS play a role in the selection of a conformation for either the canonical or non-canonical functions of NS5.

In addition to the EX1 kinetics observed in the MTase, SLA binding also has an effect on the EX1 kinetics observed in the RdRp domain. The first region in RdRp involves the fingers extension. Interestingly, such slow conformational dynamics as detected by HDX-MS have been previously reported in Hepatitis C virus NS5b, specifically the Δ2 fingers extension loop and primer grip motif which have been suggested to be regions important for the transition from initiation into the open elongation-competent state during the viral replication process [63]. One of the most interesting observation in our studies, however, is the EX1 kinetics mediated deprotection seen in the middle and pinky fingers of the RdRp domain. This deprotection is suggestive of long range allosteric effects upon SLA binding. Furthermore, it seems to be mediated by the MTase domain as the long range effects on the EX1 kinetics are largely abrogated in the absence of the MTase, showing that interdomain interactions between the MTase and RdRp domains are essential in the structure function of NS5 and its interactions with SLA. Based on its localization, this deprotection and EX1 exchange kinetics is can potentially be related to the SLA induced opening of the polymerase active site in the RdRp domain to accommodate the double-strand RNA genome. Our findings are corroborated by the thumb and fingers rearrangement during processive RNA elongation seen in the published NS5-NS3 elongation complex (EC) [22]. This has been suggested to allow RNA channeling from the NS3 helicase to the RdRp domain to perform its replicative function. However, here again, these regions also co-localize with the NS5-hSTAT2 interface. The EX1 exchange kinetics and deprotection seen in the RdRp domain regions may also reflect a conformational selection mechanism between canonical and non-canonical functions. Collectively, our studies have shed light on the different conformational rearrangements required for the different functions of MTase and RdRp domains in dengue NS5. Our cryoEM studies show that SLA binds the different dengue NS5 serotypes in a similar manner, and that this binding mechanism is most likely conserved among the various serotypes. Additionally, results from our computational modeling and HDXer studies suggest that apo DENV2 NS5 predominantly exists in the closed conformation in solution. More importantly, SLA binding imparts short-and long-range conformational effects of the EX1 kinetics type in regions that may be relevant to both of NS5’s replicative and non-canonical functions. We propose that, although predominantly in the closed conformation favored for the canonical replication functions of NS5, apo dengue NS5 can transiently sample the open conformation favored for the non-canonical functions of NS5 in a conformational selection mechanism induced by binding the relevant binding partners (Fig. 6). Ultimately, the findings from our study further add to current understanding that dengue NS5 has a complex conformational and functional landscape, which poses challenges but also provides avenues for the development of various antivirals for targeting NS5.

**Figure 6.**
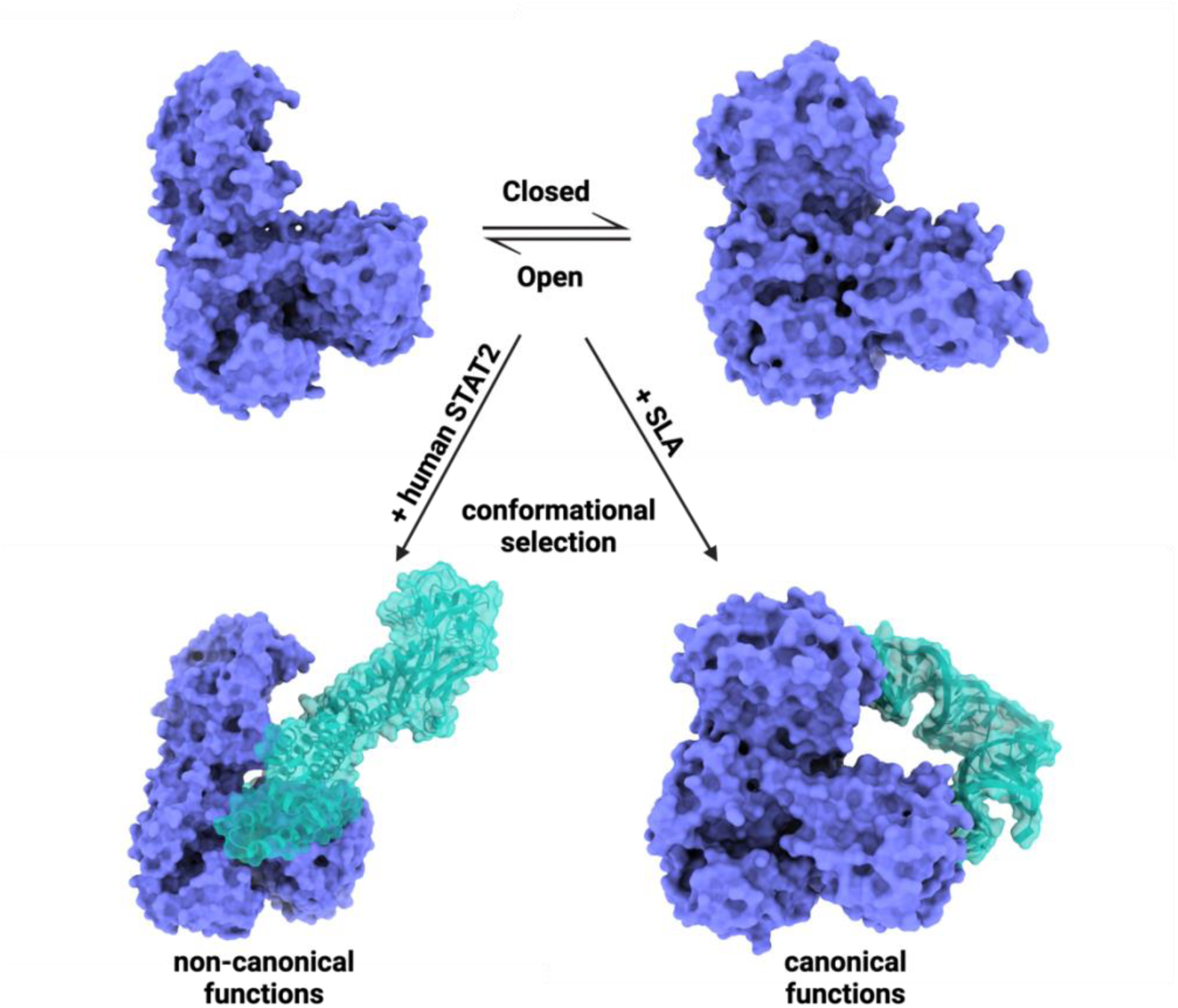
Conformational Selection Mechanism of the Dengue NS5 Protein. The open and closed conformations of dengue NS5 likely exists in equilibrium in solution. A conformational selection mechanism is proposed to occur when dengue NS5 performs its canonical functions (binding to SLA, NS3 etc.) and its non-canonical functions (binding to hSTAT2 and other host proteins).

## Author Contributions

J.O.O and D.J.D conceived the work, designed the research and developed the overall approach and methodology. K.C.K and D.J.D developed the methodology for the HDX-based refinement of docking poses and subsequent HDXer. J.O.O and G.A.S developed the cryoEM methodology. L.M performed some of the SPR experiments. J.O.O and D.J.D analyzed the data generated. J.O.O. and D.J.D. wrote the manuscript. J.O.O, K.C.K, J.K.F, G.A.S and D.J.D revised the manuscript.

## Acknowledgements

This work is supported in part by the University of Maryland, Baltimore School of Pharmacy Mass Spectrometry Center (SOP1841-IQB2014) and in part by NIH-NIGMS T32 grant GM066706 awarded to J.O.O and K.C.K. We would like to thank the Maryland Center for Advanced Molecular Analysis (M-CAMA) for microscope time and Edvin Pozharski at the University of Maryland Institute for Bioscience and Biotechnology Research (IBBR) for collecting the cryoEM data.

## Ethics declarations

### Competing Interests

The authors declare no competing interests

### Data availability

The data from this study are available from the corresponding author upon request. The cryoEM map and corresponding model have been deposited in the Electron Microscopy Data Bank (EMDB) and Protein Data Bank (PDB) with EMDB and PDB IDs of 47165 and 9DTT respectively. The HDX-MS data have been deposited to the ProteomeXchange Consortium via the PRIDE [64] partner repository with the dataset identifier PXD058297.

## Notes

### Competing Interest Statement

The authors have declared no competing interest.

