## Supplemental Data for "Structural Dynamics of the Dengue Virus Non-structural 5 (NS5) Interactions with Promoter Stem Loop A (SLA)"

^*^Corresponding author

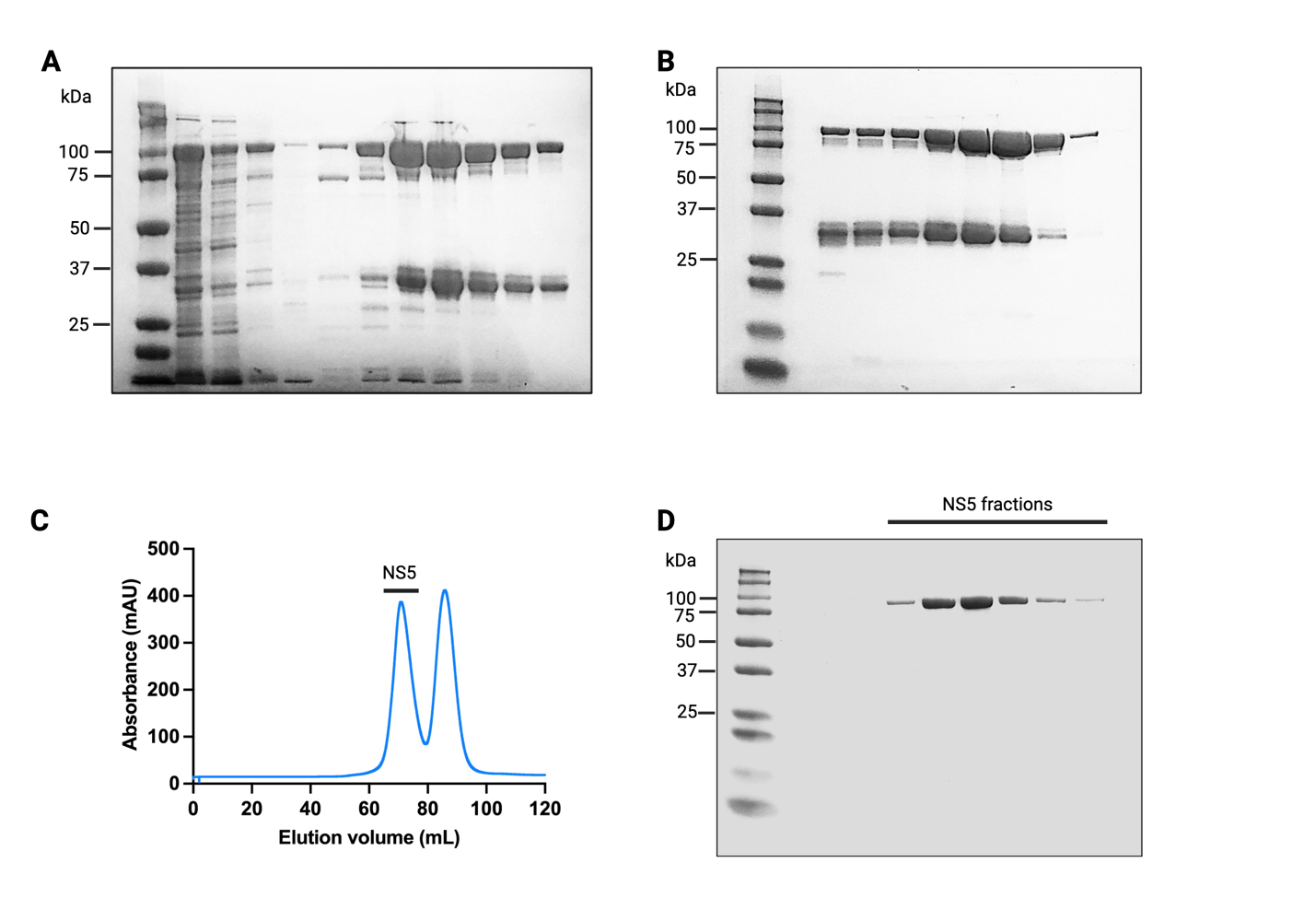

**Supplementary Figure 1. DENV2 NS5 Purification. A)** DENV2 NS5 was purified with nickel affinity purification and analyzed through SDS-PAGE. The overexpressed protein at ~105 kDa is shown with a co-purified fragment at ~32 kDa. **B)** The purified fractions from nickel affinity purification were pooled and further purified by heparin ion-exchange purification. **C and D)** The co-purified fragment was separated from DENV2 NS5 with size-exclusion chromatography.

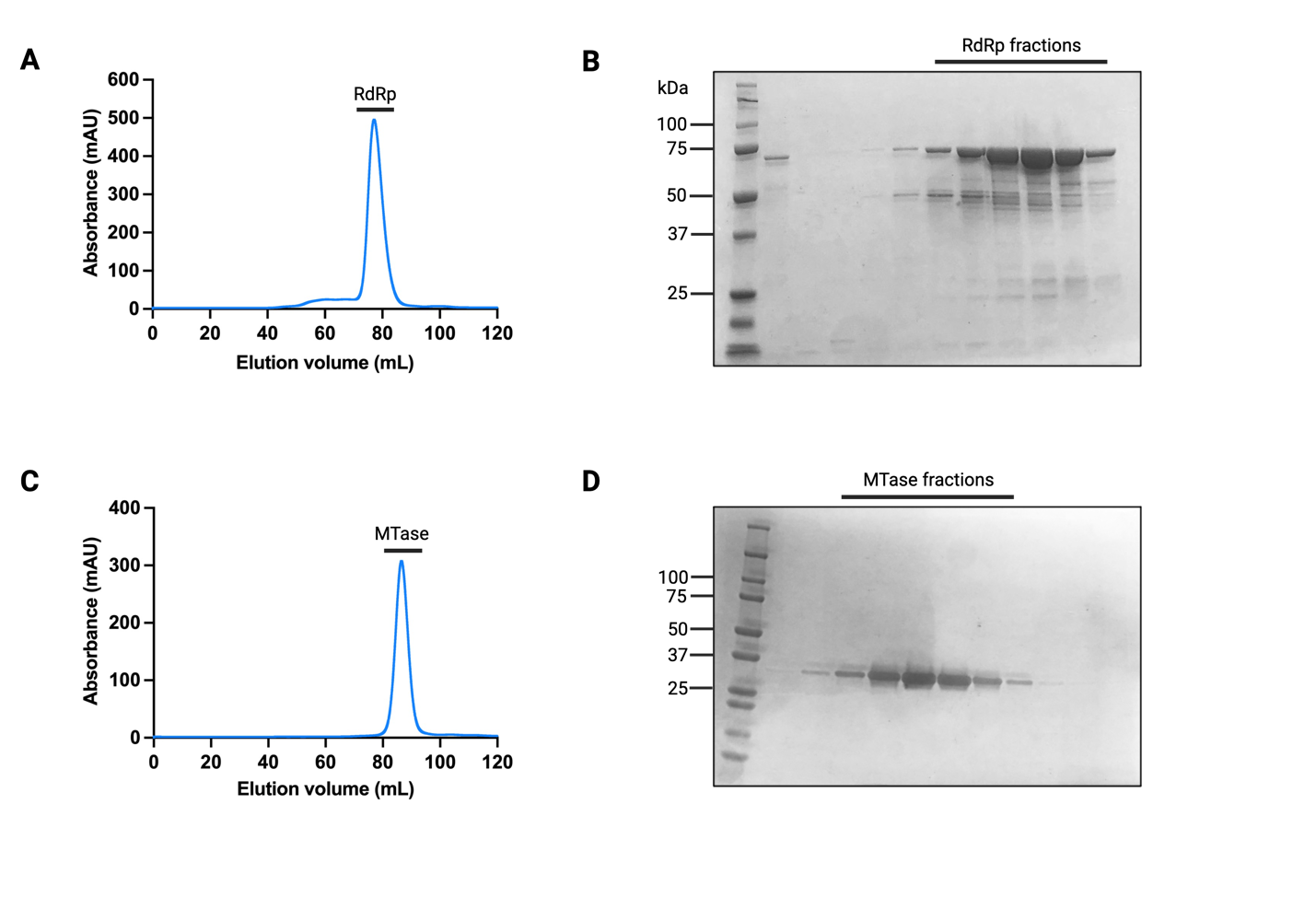

**Supplementary Figure 2. DENV2 RdRp and MTase Purification. A and B)** DENV2 NS5 RdRp purification by size-exclusion chromatography and analysis of fractions by SDS-PAGE respectively. **C and D)** DENV2 NS5 MTase purification by size-exclusion chromatography and analysis of fractions by SDS-PAGE respectively. Both RdRp and MTase domains were purified as monomers at approximately 75 kDa and 32 kDa respectively.

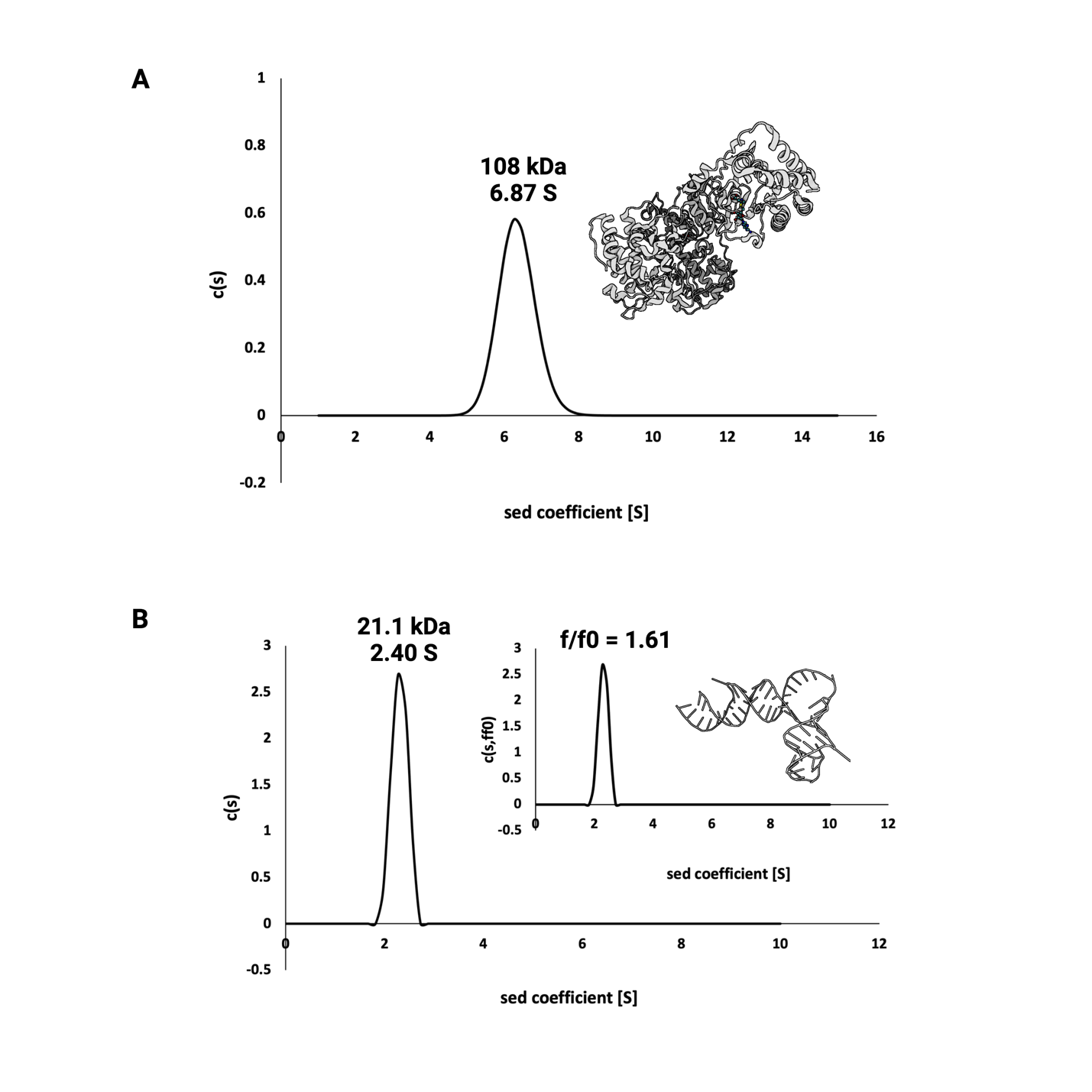
**Supplementary Figure 3. Analytical Ultracentrifugation (AUC) Sedimentation Velocity Analysis of DENV2 NS5 and SLA.** Continuous distribution plots of sedimentation coefficients are shown for DENV2 NS5 and SLA. **A)** The molecular weight of DENV2 NS5 was estimated to be ~108 kDa (actual molecular weight = 105 kDa, PDB ID shown: 5ZQK). **B)** The molecular weight of SLA was estimated to be 21.1 kDa (actual molecular weight = 22.5 kDa). The frictional ratio (f/fo) and sedimentation coefficient of SLA in 2 mM MgCl_2_ was estimated to be 1.61 and 2.40 S respectively which is similar to results obtained in previous studies by Bujalowski et al (*J Virol*, 2017). The HullRad web server was used to calculate the f/fo from the published SLA structure (PDB ID: 7LYF), with a value of 1.54 obtained.

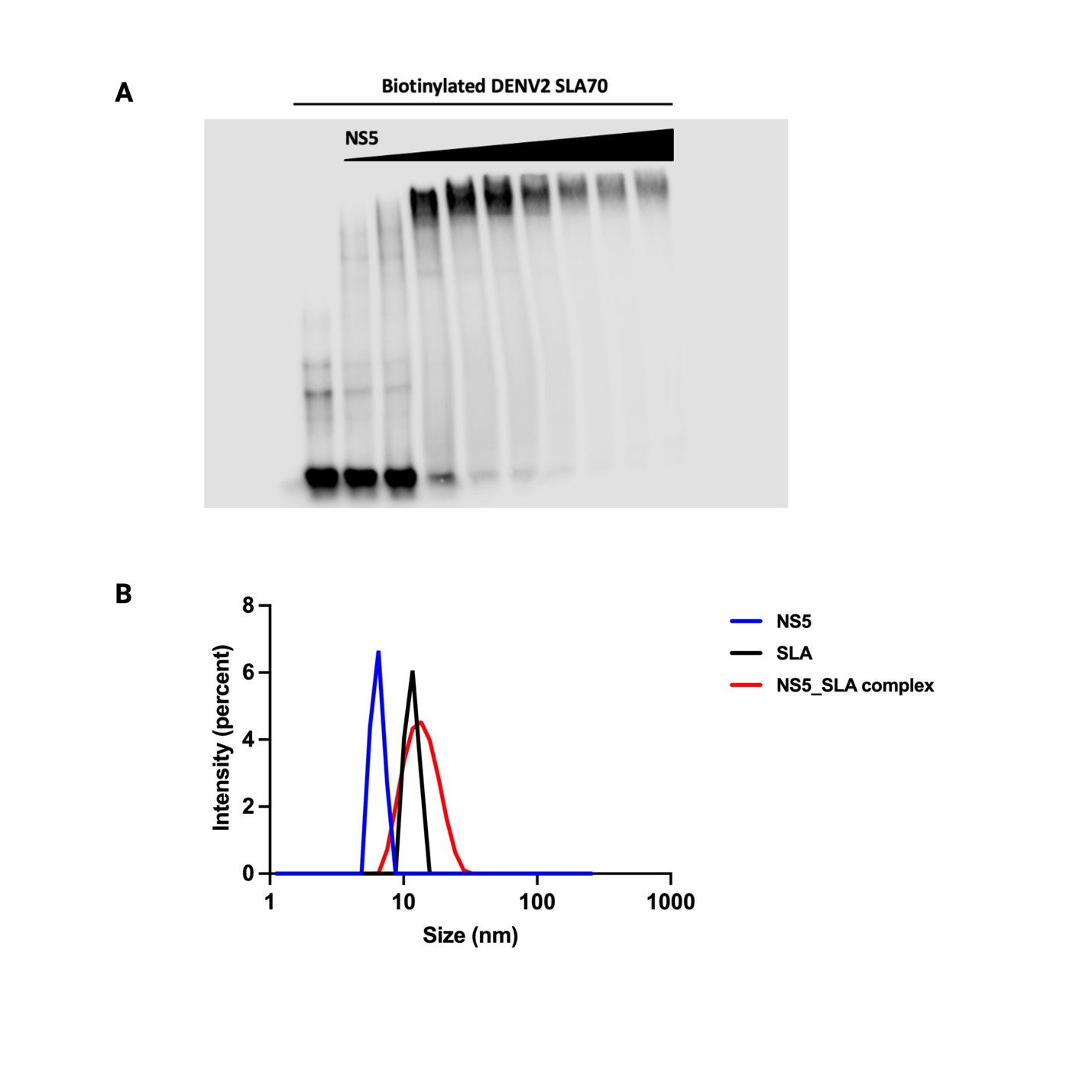
**Supplementary Figure 4. Stoichiometric Analysis of the DENV2 NS5-SLA Complex. A)** Electromobility shift (EMSA) analysis of biotin-tagged SLA saturated with DENV2 NS5. Only one band was observed in all concentrations of the NS5-SLA complex. **B)** Hydrodynamic radius analysis of particle size of individual DENV2 NS5 and SLA, and the NS5-SLA complex by dynamic light scattering.

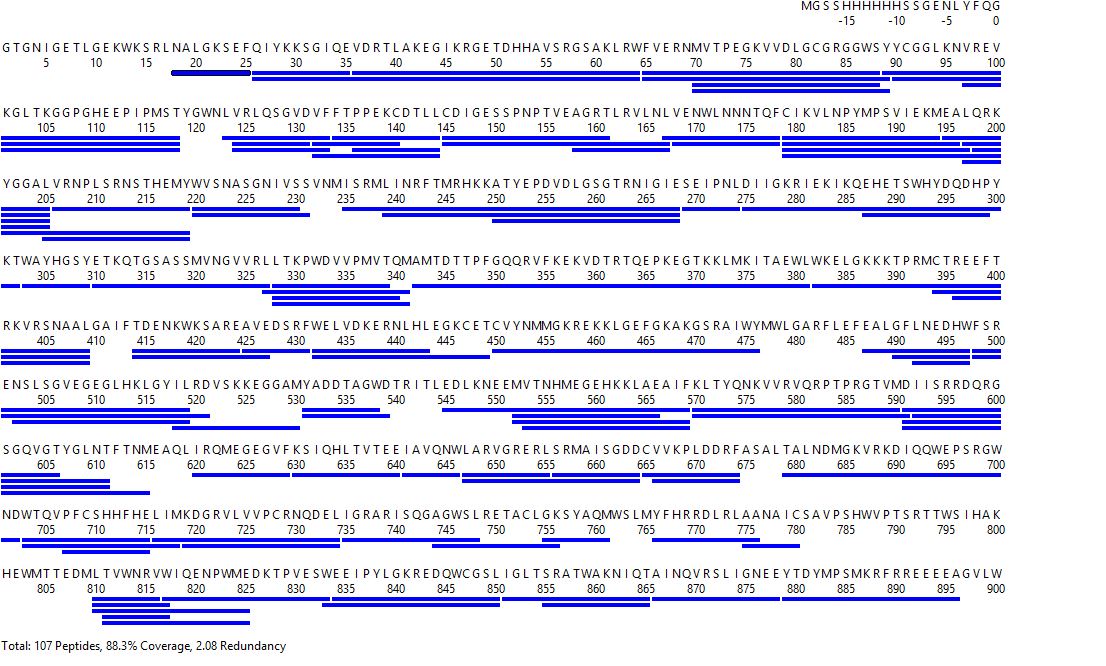

**Supplementary Figure 5. HDX-MS Sequence Coverage Map of DENV2 NS5**. Map shows peptide identification (individual peptides identified shown as blue bars) and sequence coverage analysis of undeuterated DENV2 NS5 peptic peptides showing a 88.3% sequence coverage.

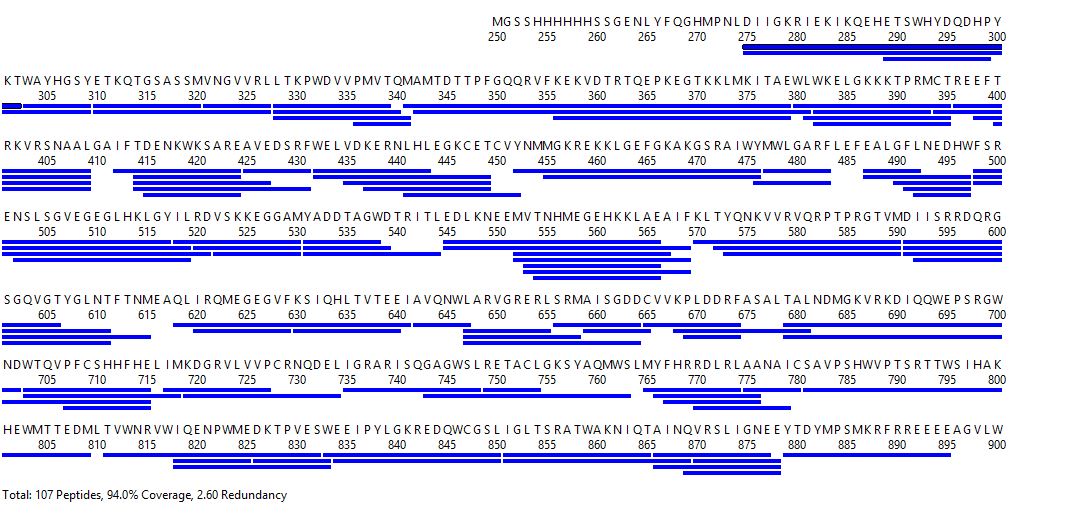

**Supplementary Figure 6. HDX-MS Sequence Coverage Map of the NS5-RdRp Domain.** Map shows peptide identification (individual peptides identified shown as blue bars) and sequence coverage analysis of undeuterated DENV2 NS5 RdRp peptic peptides showing a 94% sequence coverage.

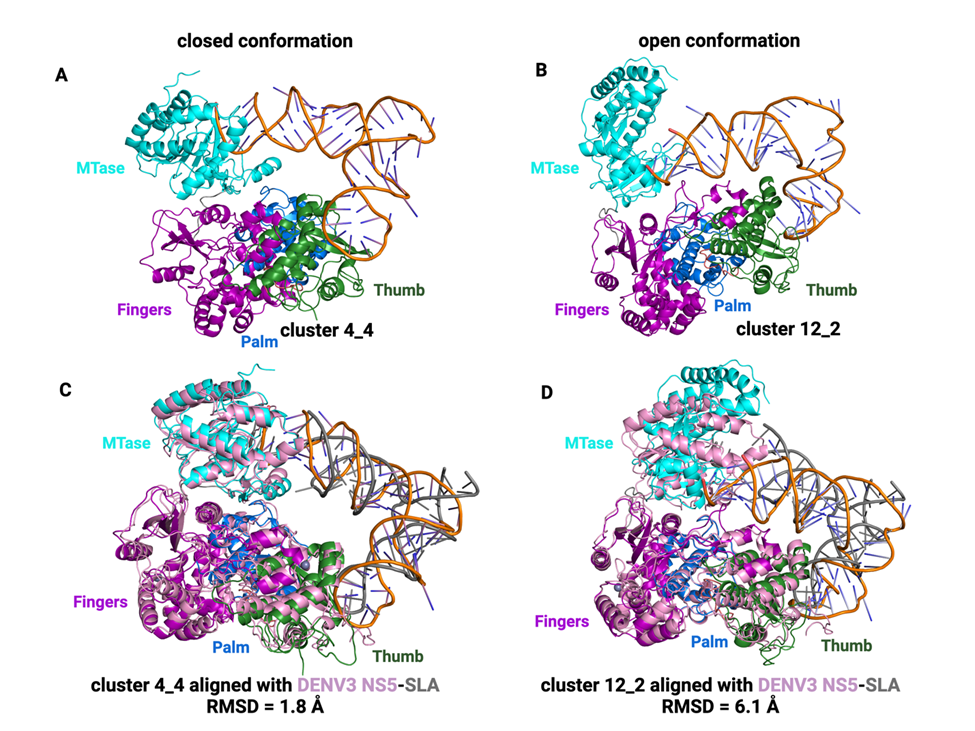

**Supplementary Figure 7. Structural Alignment of the Best-Scored Docking Poses with the DENV3 NS5-SLA CryoEM Structure.** The structures of the best-scored DENV2 NS5-SLA docking poses after applying our HDX-based scoring function from the closed and open conformation (cluster 4_4 and cluster 12_2) are shown on the top-left and top-right respectively, **A and B**. Both structures were aligned with the DENV3 NS5-SLA cryoEM structure (PDB ID 8GZP) with RMSD values of 1.80 Å and 6.08 Å respectively as shown on the bottom-left and bottom-right, **C and D**.

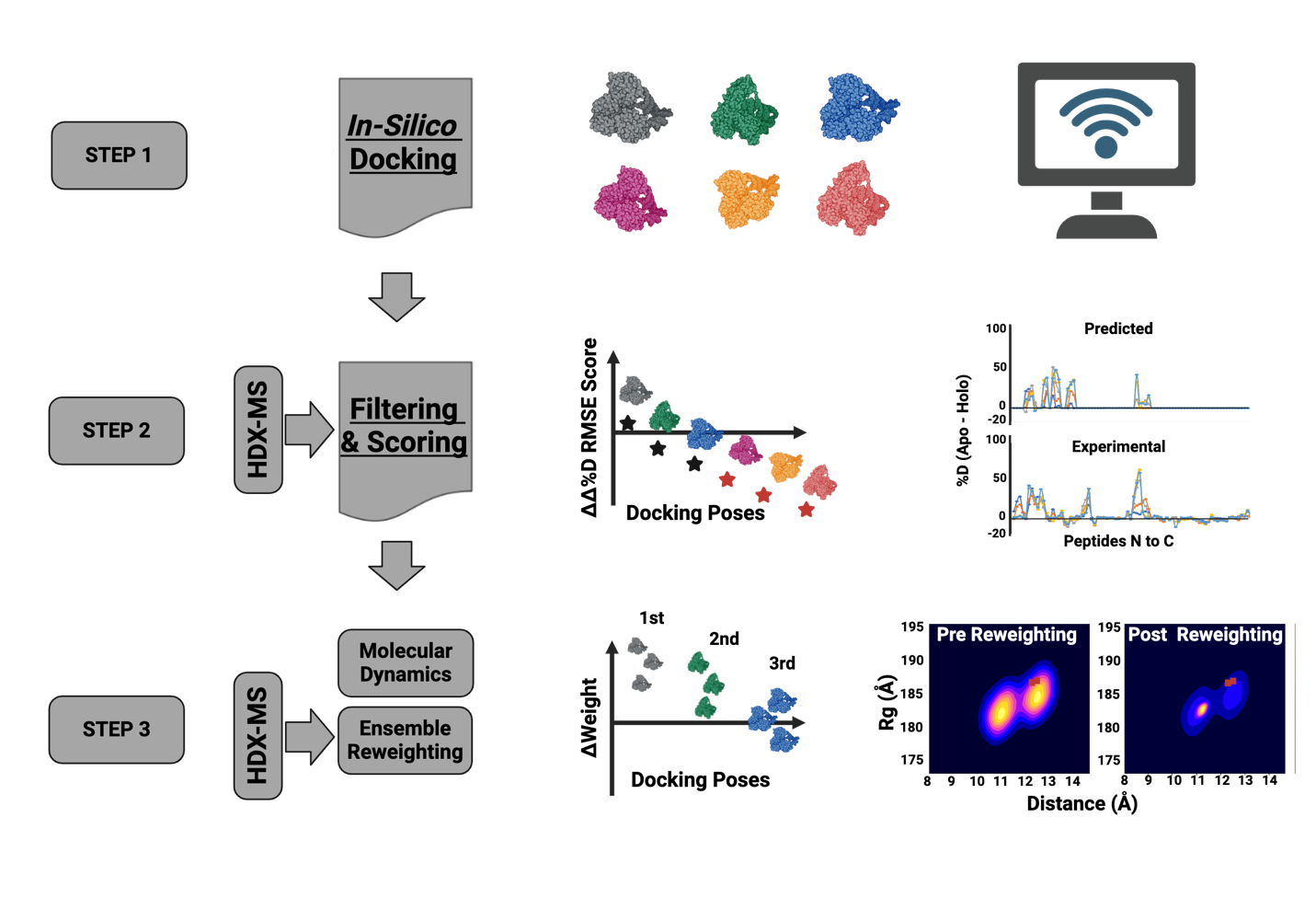

**Supplementary Figure 8. Summary of the Hydrogen-Deuterium Exchange Ensemble Reweighting (HDXer) Workflow.** The workflow includes in-silico docking using the HADDOCK web server in step 1, followed by HDX-based filtering and scoring of the HADDOCK-generated poses in step 2, and lastly MD simulations and HDXer in step 3 to determine the populations of frames that conform the most with the experimental HDX-MS data.

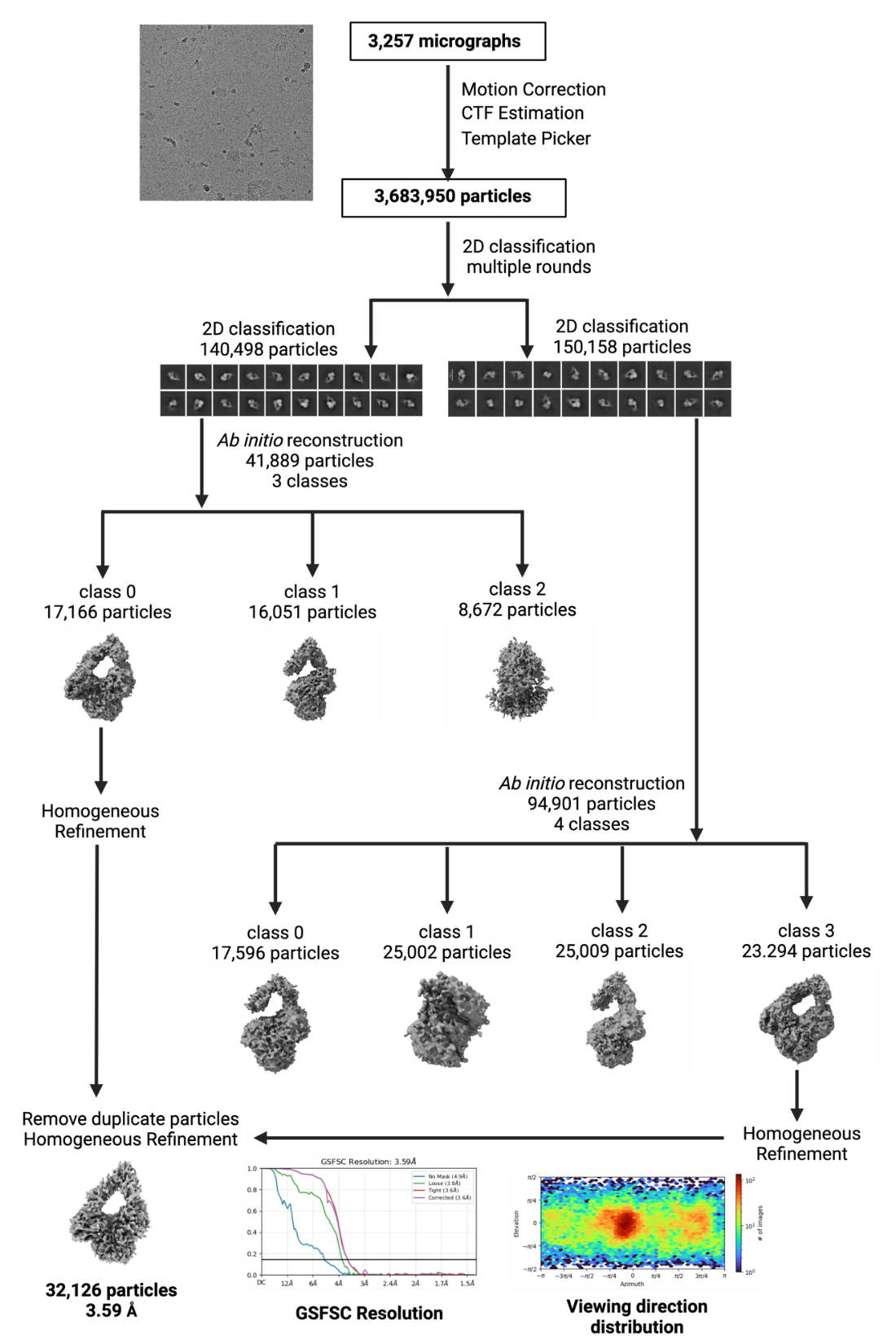

**Supplementary Figure 9. Summary of the cryoEM processing workflow for the DENV2 NS5-SLA complex (EMD-47165, PDB ID: 9DTT).** Data processing for cryoEM reconstruction and final map was done in cryoSPARC. A final map with 3.59 Å from 32,126 particles was used for building the final model deposited in the PDB.

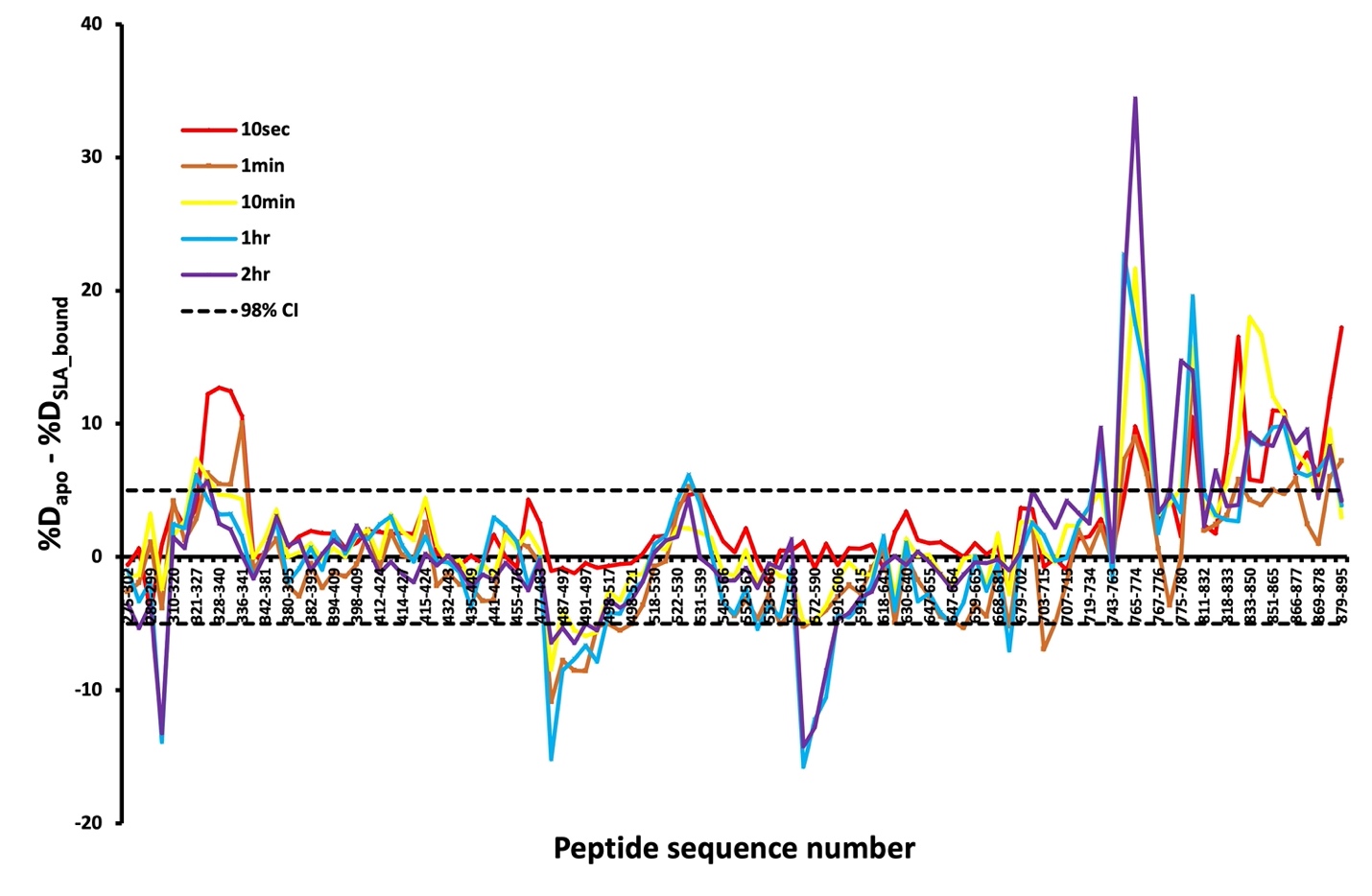

**Supplementary Figure 10. Differences in Percent Deuterium Incorporation of Apo Versus SLA-bound NS5-RdRp Domain.** Peptide fragments from N- to C- terminal of the RdRp domain alone are shown from left to right based on the residue number of the first amino acid of each peptide. The difference in percent deuterium incorporation between the apo and SLA-bound states (Δ%D) is plotted for each peptide fragment and for each deuteration time point. The color coding used is based on the deuteration time points probed. The horizontal dashed lines indicate 98% confidence intervals.

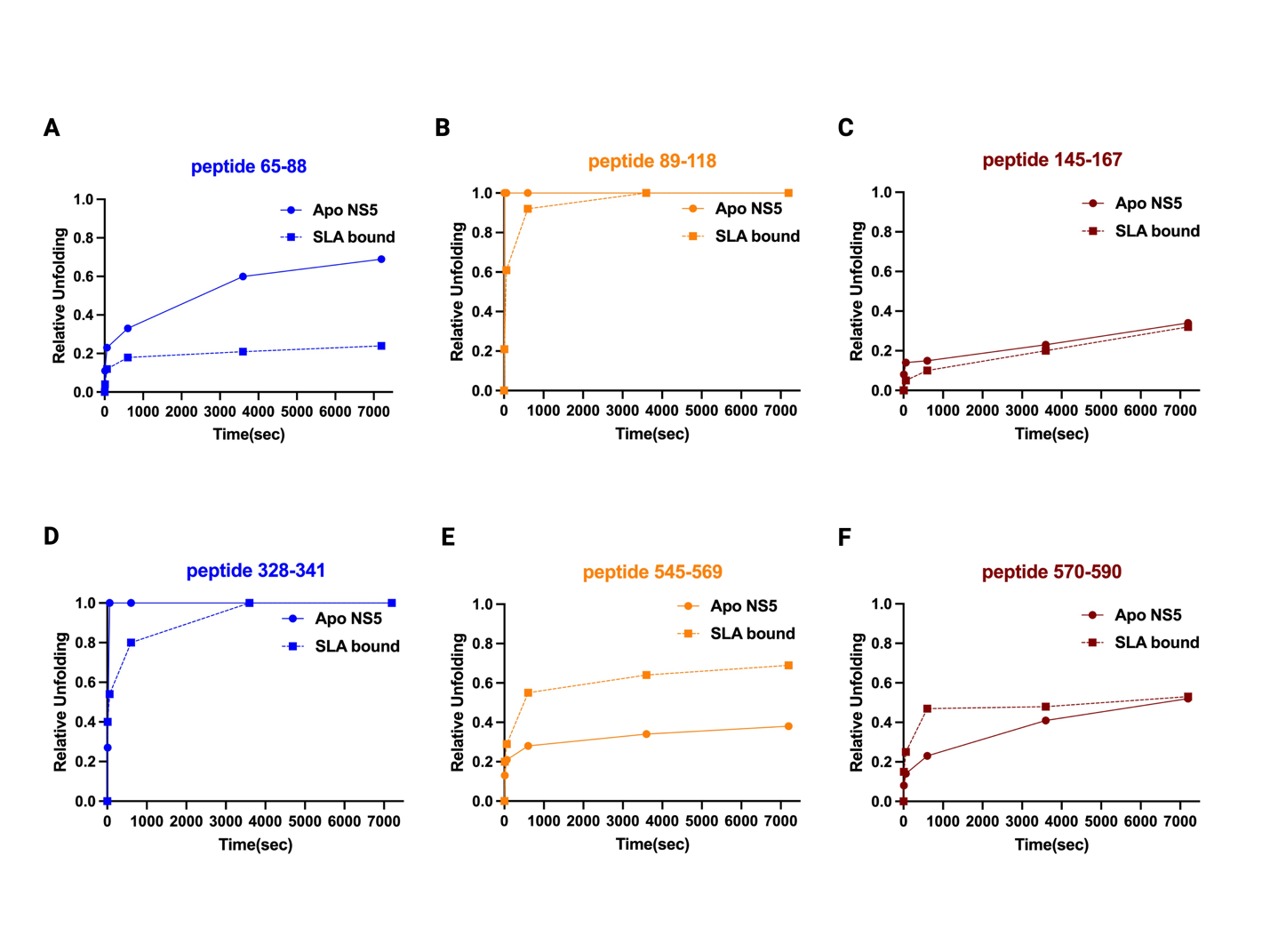

**Supplementary Figure 11. Modulation of the EX1 Exchange kinetics in Full-length DENV2 NS5 upon SLA Binding.** The relative appearance of the highly deuterated species for all EX1 peptides at time points 0 s, 10 s, 60 s, 3600 s and 7200 s are shown. The relative appearance of the highly deuterated species in EX1 peptides 65-88, 89-118 and 145-167 in the MTase domain is generally decreased in the presence of SLA **(A-C)**. The greatest decrease in the presence of SLA is observed in peptide 65-88. The relative appearance of the highly deuterated species for EX1 peptides in the RdRp domain is decreased for peptide 328-341, and increased for peptides 545-549 and 570-590 respectively, with peptide 545-549 showing a greater increase in the appearance of the highly deuterated species in the presence of SLA **(D-F)**.

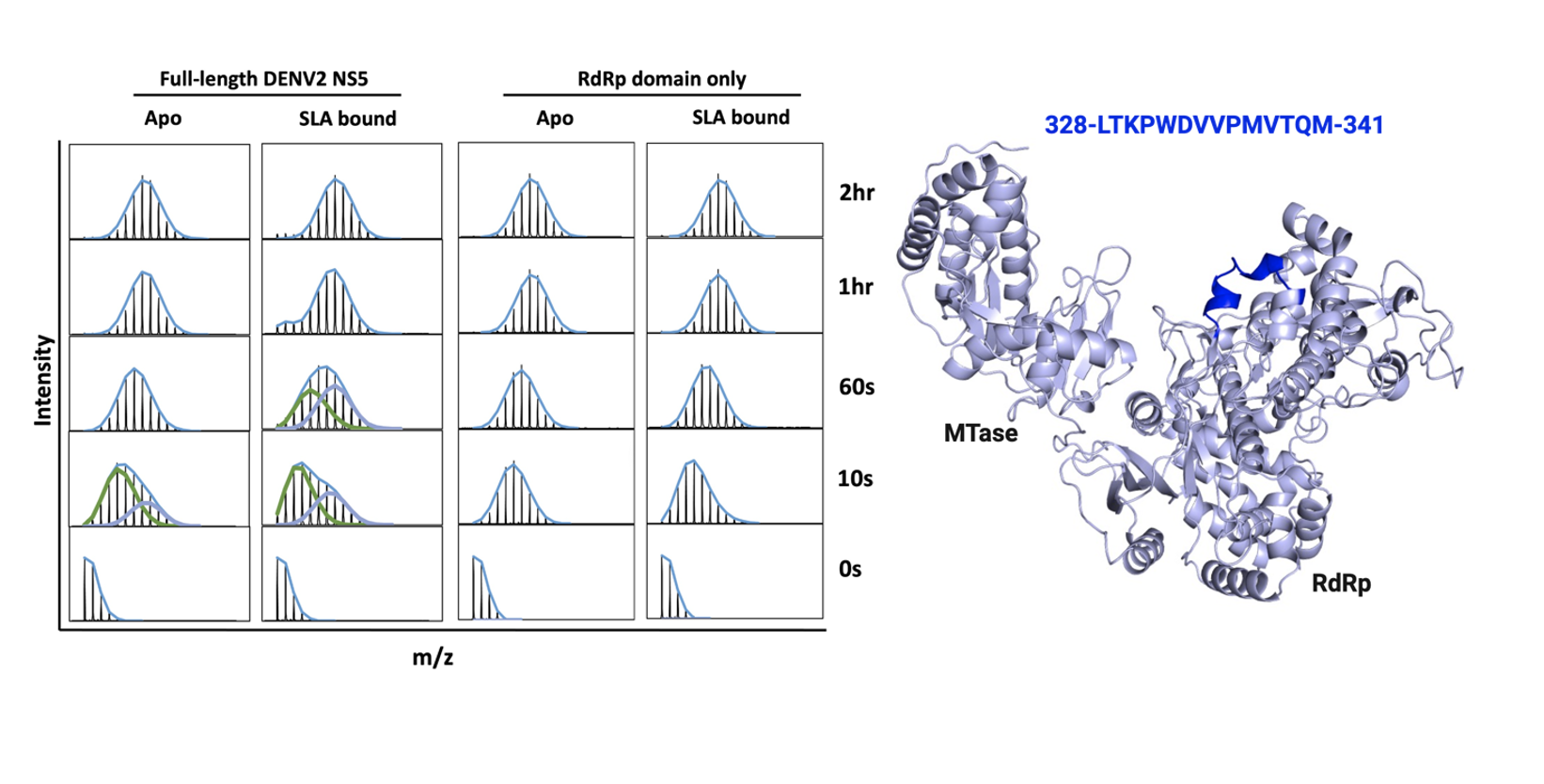

**Supplementary Figure 12. Deconvolution of Bimodal Isotopic Envelopes for Peptide 328-341 in the NS5-RdRp Domain.** The deconvoluted bimodal isotopic envelopes are shown in the apo form and SLA-bound form in the context of full-length DENV2 NS5 and the NS5-RdRp domain alone (left). While bimodal isotopic envelopes were observed in peptide 328-341 in the RdRp domain for full-length NS5, no EX1 bimodal behavior was observed for the NS5 RdRp domain only (both apo and SLA-bound). Peptide 328-341 is located in the index finger of the NS5-RdRp domain which extends into the thumb sub-domain (right).

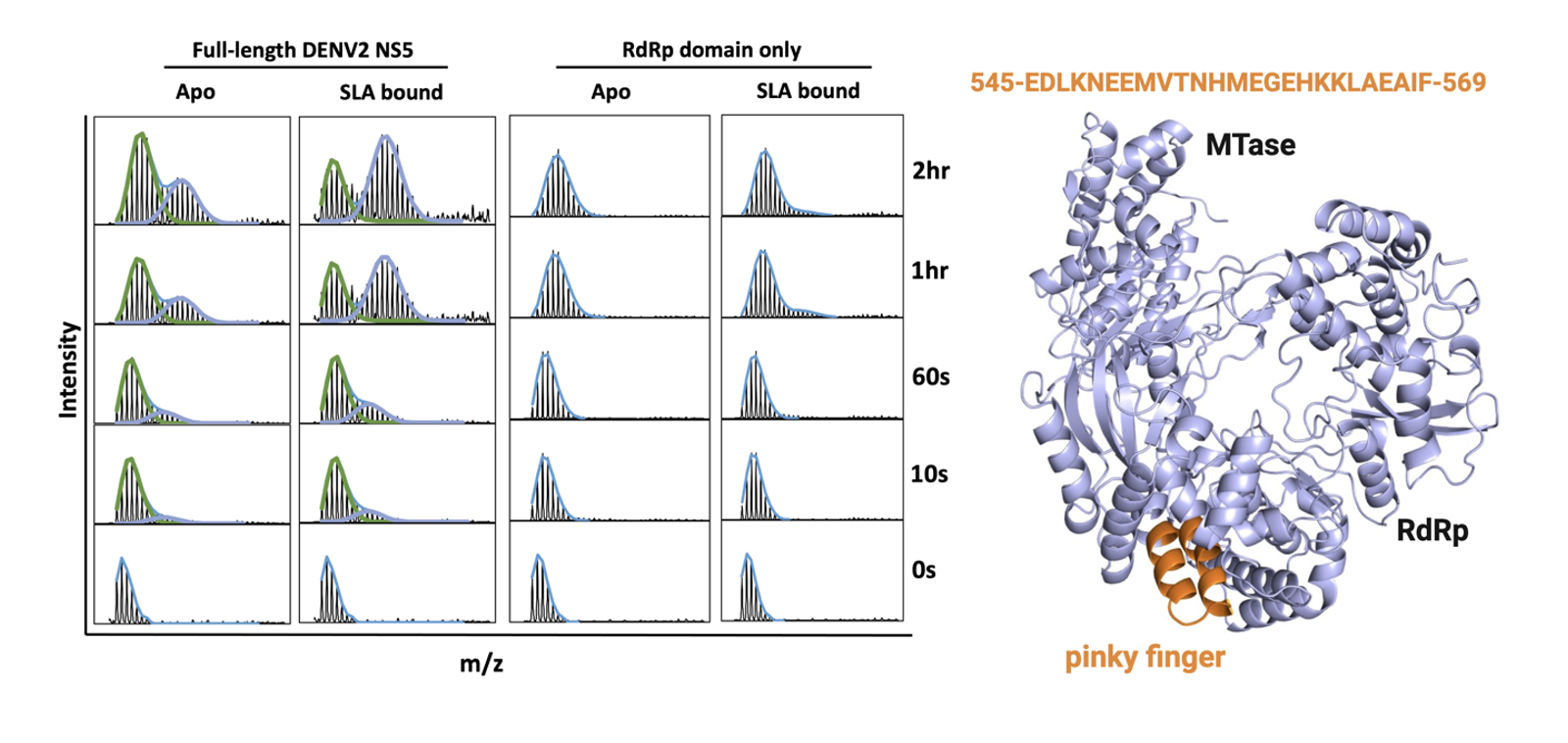

**Supplementary Figure 13. Deconvolution of Bimodal Isotopic Envelopes for Peptide 545-569 in the NS5-RdRp Domain.** The deconvoluted bimodal isotopic envelopes are shown in the apo form and SLA-bound form in the context of full-length DENV2 NS5 and the NS5-RdRp domain alone (left). No EX1 bimodal behavior was observed for the NS5 RdRp domain only (both apo and SLA-bound). Peptide 545-569 is located at the pinky finger of the NS5-RdRp in the fingers sub-domain (right).

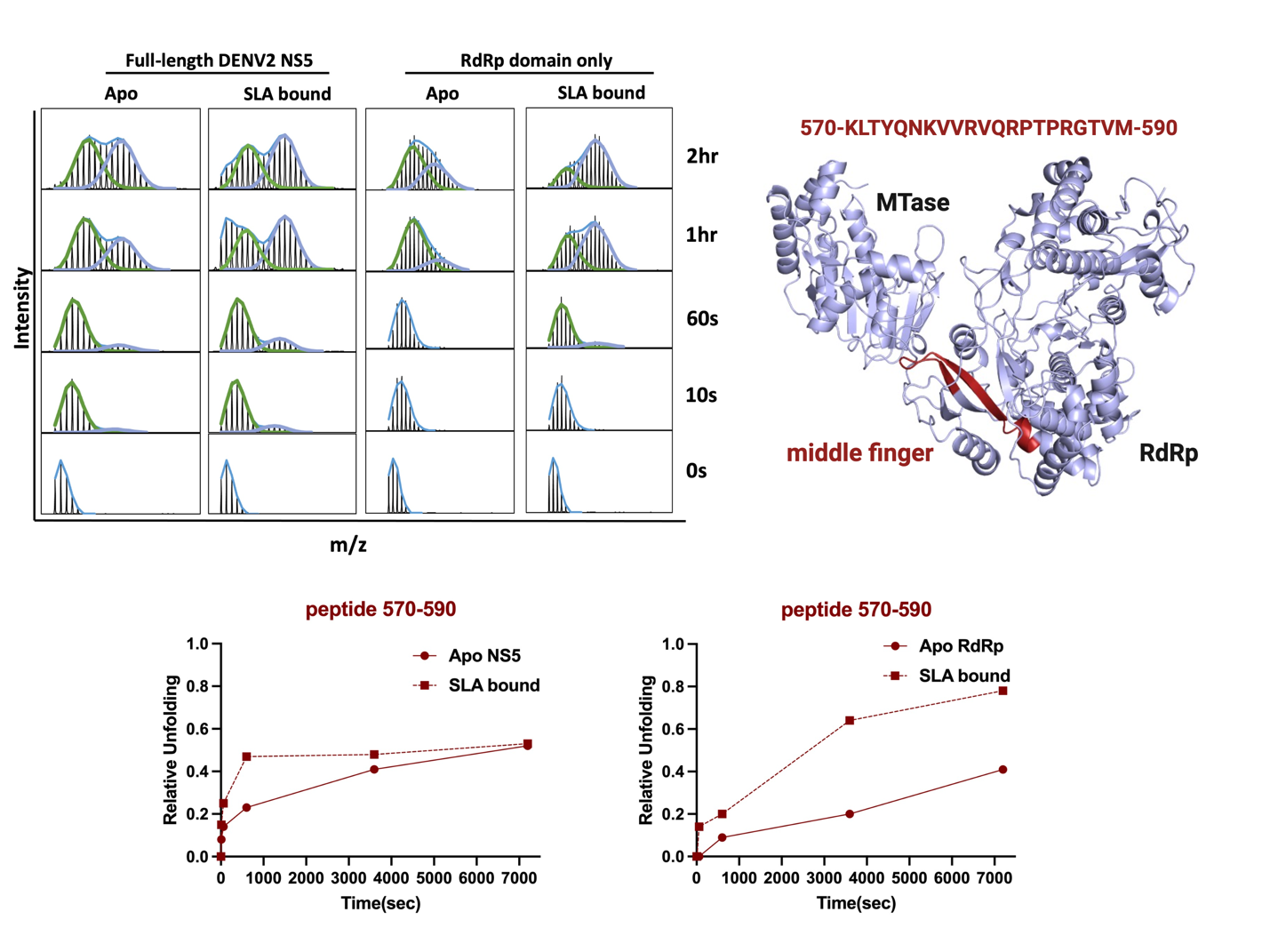

**Supplementary Figure 14. Deconvolution of Bimodal Isotopic Envelopes for Peptide 570-590 in the NS5-RdRp Domain.** The deconvoluted bimodal isotopic envelopes are shown in the apo form and SLA-bound form in the context of full-length DENV2 NS5 and the NS5-RdRp domain alone (upper left). EX1 bimodal behavior was observed for this peptide in both full-length NS5 RdRp and the NS5-RdRp domain only (both apo and SLA-bound). In both cases, the rate of appearance of the highly deuterated species was increased in the presence of SLA, with a much higher effect seen in the context of the NS5-RdRp domain only (lower panel). Peptide 545-569 is located at the middle finger of the RdRp in the fingers sub-domain (upper right).

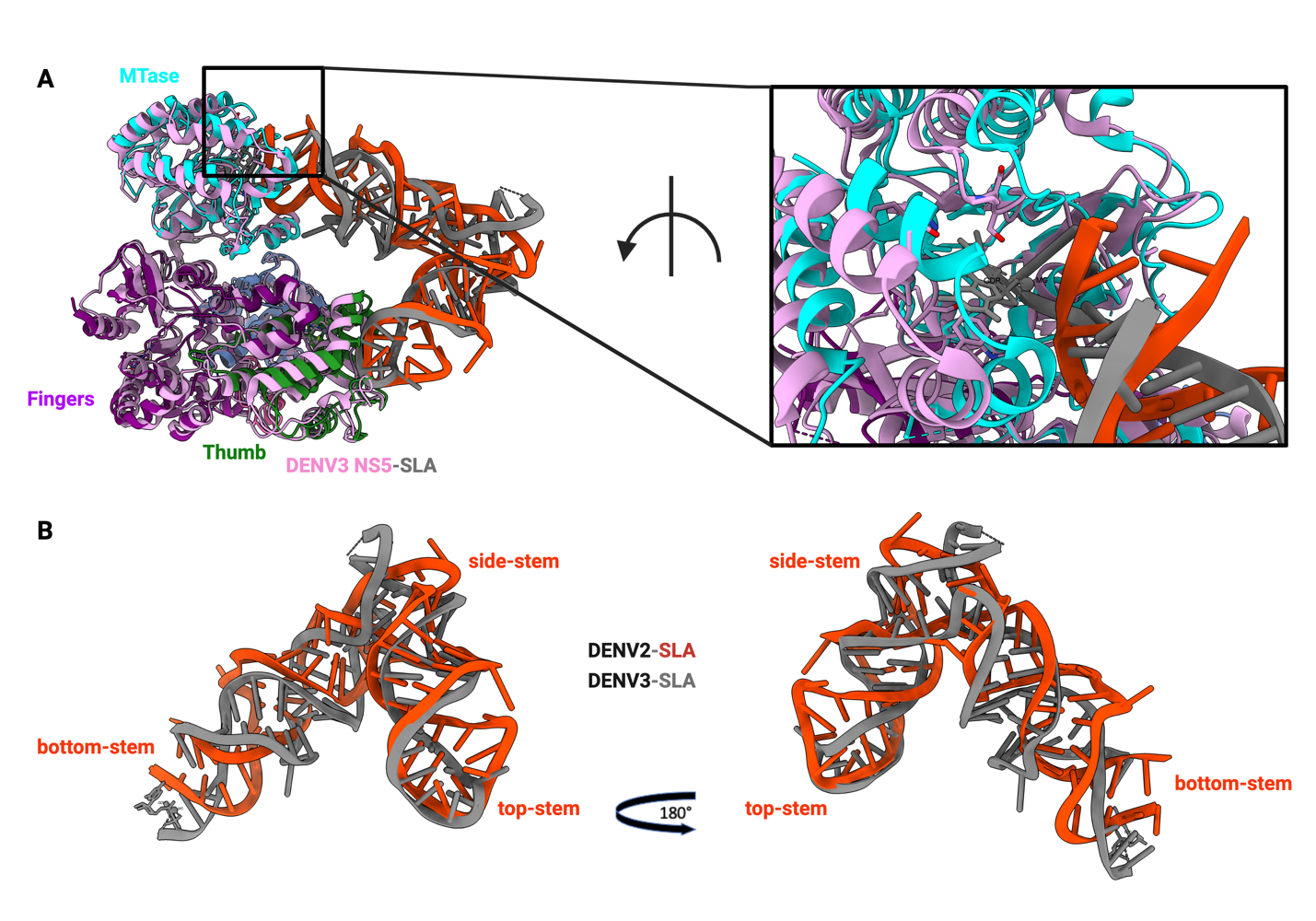

**Supplementary Figure 15. Structural Alignment of the DENV2 and DENV3 NS5-SLA CryoEM Structures. A.** Alignment of the cryoEM NS5-SLA complex from DENV2 (our work) and DENV3 (PDB: 8GZP) gave an RMSD of 2.16 Å. The SLA bottom stem loop binding in the MTase catalytic site from both complexes are shown in the enlarged figure (top right panel) **B.** Alignment of SLA from both DENV2 and DENV3 cryoEM NS5-SLA complexes, showing the bottom-stem, side-stem, and top-stem loops respectively.

**Supplementary Table 1. HDX Data Summary.**

| **Data Set** | **Apo DENV2 NS5** | **NS5-SLA** |
| --- | --- | --- |
| **HDX reaction details** | 80% D20, pD 7.0 (pH 7.4), 25ºC | 80% D20, pD 7.0 (pH 7.4), 25ºC |
| **HDX time course (min)** | 0, 0.167, 1, 10, 60, 120 | 0, 0.167, 1, 10, 60, 120 |
| **HDX controls** | undeuterated control; fully deuterated control | undeuterated control; fully deuterated control |
| **Back-exchange (%)** | 20.6 (IQR = 17.4) | 24.9 (IQR = 19.2) |
| **Number of Peptides** | 107 | 107 |
| **Sequence coverage** | 88% | 88% |
| **Average peptide (Length / Redundancy** | 14.12 / 2.08 | 14.12 / 2.08 |
| **Replicates** | 3 (technical) | 3 (technical) |
| **Repeatability (average standard deviation)** | 0.069 | 0.086 |
| **Significant differences in HDX (Δ%D)** | N/A | 98% CI = 7.08 (Apo minus SLA bound) |
| **Data Set** | **Apo NS5-RdRp** | **RdRp-SLA** |
| **HDX reaction details** | 80% D20, pD 7.0 (pH 7.4), 25ºC | 80% D20, pD 7.0 (pH 7.4), 25ºC |
| **HDX time course (min)** | 0, 0.167, 1, 10, 60, 120 | 0, 0.167, 1, 10, 60, 120 |
| **HDX controls** | undeuterated control; fully deuterated control | undeuterated control; fully deuterated control |
| **Back-exchange (%)** | 18.3 (IQR = 14.7) | 17.2 (IQR = 13.8) |
| **Number of Peptides** | 107 | 107 |
| **Sequence coverage** | 94% | 94% |
| **Average peptide (Length / Redundancy** | 13.43 / 2.60 | 13.43 / 2.60 |
| **Replicates** | 3 (technical) | 3 (technical) |
| **Repeatability (average standard deviation)** | 0.058 | 0.056 |
| **Significant differences in HDX (Δ%D)** | N/A | 98% CI = 5.82 (Apo minus SLA bound) |

**Supplementary Table 2. Cryo-EM data collection, refinement and validation statistics.**

|  | **DENV2 NS5-SLA Complex**  **(EMD-47165)**  **(PDB 9DTT)** |
| --- | --- |
| **Data collection and processing** |  |
| Microscope  Camera  Magnification | Talos Arctica  Gatan K3  70,000 |
| Voltage (kV) | 200 |
| Electron exposure (e–/Å^2^) | 46.077 |
| Defocus range (μm) | -1.1 to -2.0 |
| Pixel size (Å)  Micrographs (no.) | 0.718  3,257 |
| Initial particles (no.) | 3,683,950 |
| Symmetry imposed | C1 |
| Initial particle images (no.) | 290,656 |
| Final particle images (no.)  Map resolution (Å) | 32,126  3.59 |
| FSC threshold | 0.143 |
| **Refinement** |  |
| Initial model used | 6KR3, 7LYF |
| Model resolution (Å) | 3.59 |
| FSC threshold  Map sharpening *B* factor (Å^2^) | 0.143  -70 |
| Model composition |  |
| Non-hydrogen atoms | 8,274 |
| Protein residues  Nucleotide residues  Ligands (zinc)  *B* factors (Å^2^) | 850  67  2 |
| Protein  RNA  Ligand | 77.82  235.15  82.84 |
| R.m.s. deviations  Bond lengths (Å)  Bond angles (°) | 0.002  0.486 |
| **Validation**  MolProbity score  Clashscore  Poor rotamers (%) | 1.87  10.36  0 |
| Ramachandran plot  Favored (%)  Allowed (%)  Disallowed (%) | 95.18  4.82  0 |

**Supplementary Table 3. HDX-based scoring of HADDOCK-generated DENV2 NS5-SLA docking poses suggest SLA binding does not induce opening or closing of NS5.** Δ%D RMSE of the DENV2 NS5-SLA docking poses in the closed confirmation versus the apo DENV2 NS5 in the closed or open conformation (Left) and Δ%D RMSE of the DENV2 NS5-SLA docking poses in the open confirmation versus the apo DENV2 NS5 in the closed or open conformation (Right) show that SLA binding is not associated with conversion from open to close, or from close to open conformations.

|  | **Closed SLA-bound** |  |  | **Open SLA-Bound** |  |
| --- | --- | --- | --- | --- | --- |
| **Model** | **Δ%D RMSE (Apo, closed)** | **Δ%D RMSE (Apo, open)** | **Model** | **Δ%D RMSE (Apo, open)** | **Δ%D RMSE (Apo, closed)** |
| Cluster 1_1 | 0.1333 | 0.1700 | Cluster 3_1 | 0.1480 | 0.1772 |
| Cluster 1_2 | 0.1387 | 0.1727 | Cluster 3_2 | 0.1460 | 0.1762 |
| Cluster 1_3 | 0.1394 | 0.1732 | Cluster 3_3 | 0.1499 | 0.1745 |
| Cluster 1_4 | 0.1369 | 0.1727 | Cluster 3_4 | 0.1437 | 0.1772 |
| Cluster 4_1 | 0.1365 | 0.1908 | Cluster 8_1 | 0.1483 | 0.1818 |
| Cluster 4_2 | 0.1389 | 0.1820 | Cluster 8_2 | 0.1452 | 0.1768 |
| Cluster 4_3 | 0.1392 | 0.1855 | Cluster 8_3 | 0.1451 | 0.1782 |
| Cluster 4_4 | 0.1265 | 0.1780 | Cluster 8_4 | 0.1465 | 0.1764 |
